# Physics-Guided Neural Reconstruction of Cellular Membranes for 3D Electron Microscopy

**DOI:** 10.64898/2026.08.01.742159

**Authors:** Atsushi Matsuda, Se Min Kim, Matthew Akamatsu, Christopher T. Lee

**Affiliations:** Department of Biology, University of Washington; Department of Molecular Biology, University of California San Diego

## Abstract

With advances in three-dimensional electron microscopy modalities, quantitative characterization of membrane ultrastructure has emerged as an approach to interrogate how organization of proteins and other components around the membrane drive structure and function. Hindering these efforts, the confident reconstruction of geometric features such as membrane curvature is challenging since it requires the calculation of higher-order derivatives from discrete membrane representations. Mod-ern advances in using neural networks to learn continuous implicit representations of complex shapes present a promising solution to this problem. This work presents a physics-informed neural network framework for reconstructing membrane geometries to curvature-order accuracy from images using an implicit neural representation. Benchmarking using synthetic data illustrates that physics-based regularization during training improves accuracy of recovered curvatures, improving robustness to image noise. Applications to experimental datasets demonstrate that the framework generalizes to complex cellular membrane structures, such as the Golgi apparatus and mitochondria. We further apply the framework to endoplasmic reticulum–plasma membrane contact sites, demonstrating its ability to capture localized membrane deformations associated with membrane–membrane interactions. We also perform three-dimensional curvature analysis of endocytic pits in cells, providing preliminary evidence of anisotropic curvature patterns at the pit neck consistent with theoretical predictions of a lower-energy pathway for neck constriction.

## B Introduction

Biological membranes define the structural and functional organization of cells, establishing compartmental boundaries that regulate biochemical processes and communication across cellular environments.^1,2^ Far from being passive barriers, membranes are dynamic interfaces that reshape to support processes such as endocytosis, exocytosis, intracellular trafficking and protein sorting, cell migration, and organelle biogenesis. Through these deformations, membranes provide a plat-form where molecular interactions are translated into large-scale structural changes, enabling cells to adapt to mechanical and biochemical cues. As such, membrane morphology plays a central role in cellular function, making quantitative characterization of membrane geometry essential for understanding the underlying mechanisms of membrane-associated biological processes.

Among geometric descriptors of membrane shape, curvature occupies a central role because it links observable membrane geometry to the molecular and mechanical processes that govern membrane remodeling.^3,4^ Spatial variations in membrane curvature reciprocally influence the positioning of curvature-sensitive proteins as well as the mechanical forces acting on the membrane.^5^ Consequently, quantitative measurements of membrane curvature provide insights that extend be-yond membrane morphology alone, enabling investigation of the molecular and mechanical basis of cellular processes.^2,6^ For example, membrane deformations have been used to infer the membrane footprint generated by mechanosensitive proteins such as Piezo,^7^ while curvature-based analyses combined with membrane elasticity theory have enabled inference of physical forces and material properties from membrane shape itself.^8–11^ These studies demonstrate that membrane geometry serves as a quantitative reporter of molecular interactions and mechanical forces that are otherwise difficult to measure directly.

Realizing the full potential of curvature-based analysis requires accurate three-dimensional reconstruction of membrane geometry from volumetric imaging data. Advances in three-dimensional electron microscopy, including cryo-electron tomography and volume electron microscopy (vEM) techniques such as focused ion beam scanning electron microscopy (FIB-SEM), now enable visualization of membrane ultrastructure at nanometer resolution within intact cells.^12–15^ In parallel, recent advances in machine-learning-based membrane segmentation, such as MemBrain-seg, TARDIS, TomoSegNet, and ETSAM, have substantially improved automated reconstruction of membrane structures from volumetric imaging data.^16–19^ Together, these technological advances have enabled quantitative analysis of membrane geometry at an unprecedented scale and resolution.

Despite these advances, quantitative membrane-mechanics analyses have been largely limited to simple membrane systems, such as synthetic vesicles.^7,11,20^ Extending these analyses to biologically relevant complex cellular membranes remains challenging because biological membranes exhibit heterogeneous morphologies, intricate topology, and are imaged under inherently noisy conditions. A key challenge lies in accurately estimating geometric properties from reconstructed membrane surfaces. Most computational pipelines represent membranes using discrete formats such as voxel segmentations or triangulated meshes.^21–23^ Although these representations provide established frameworks for curvature estimation, the accuracy of such estimates can be sensitive to spatial resolution, mesh quality, and noise in the reconstructed geometry. Consequently, small geometric errors introduced during segmentation or meshing are amplified during numerical differentiation, often resulting in inaccurate curvature estimates and limiting downstream quantitative membrane-mechanics analyses (Fig. 1A).

**Figure 1:**
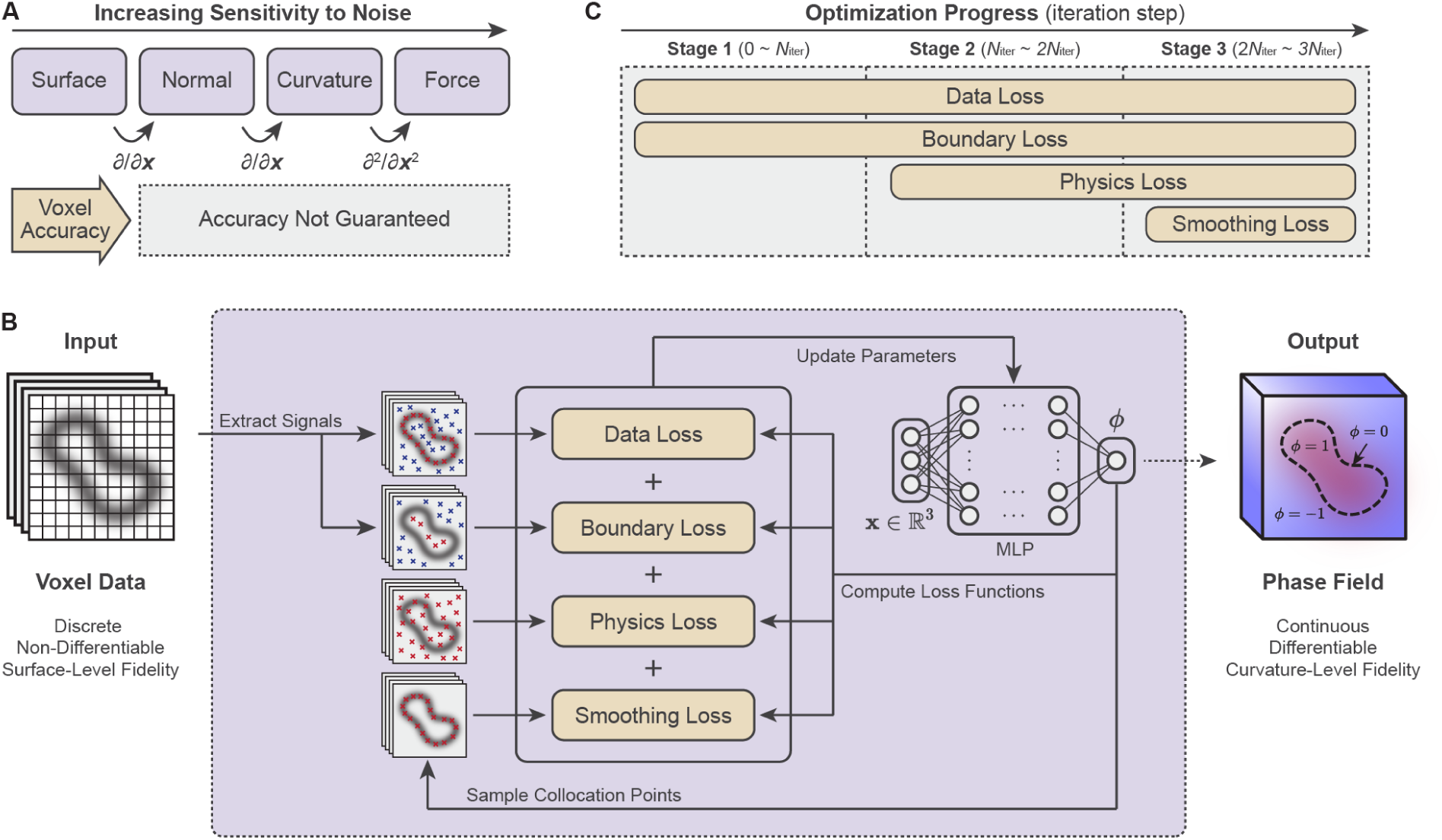
Our framework combines implicit neural representations (INRs) with physics-informed neural networks (PINNs) to reconstruct membrane geometry from volumetric imaging data with curvature-level accuracy. (A) Differential geometric quantities are calculated by taking higher-order derivatives of the surface representation. Accurate calculation of differential geometric quantities requires progressively higher-order derivatives of the surface representation, through which reconstruction errors can accumulate and propagate. (B) We aim to reconstruct membrane geometry from volumetric imaging data using a phase-field representation. In the phase field, the membrane is represented as a continuous scalar field, where values of *φ* = ±1 correspond to the bulk regions and the zero level set defines the membrane surface. The field is parameterized by an implicit neural representation trained via multi-objective optimization to balance fit to image data and consistency with physics. (C) Optimization is performed across three sequential stages to stabilize training. At each stage, additional objectives are added to refine the solution.

A natural solution to this challenge is to represent membrane geometry as a continuous differentiable function rather than a discrete surface. Implicit neural representations (INRs)^24,25^ use a fully connected neural network to represent a continuous scalar field over the spatial domain; they are functions which take spatial coordinates as input, outputting a scalar field whose level set defines the underlying geometry. By representing the geometry via an implicit function encoded by a neural network, INRs are both smooth and differentiable, which addresses the challenges of working with discrete representations and allowing relevant geometric quantities such as curvature to be computed directly.

Building on this idea, we recast membrane shape reconstruction as the problem of learning an INR, constrained by the volumetric imaging data, from which the geometry and curvature can be quantified (Fig. 1B). Although the image data provide information about membrane location, they do not constrain curvature, making the problem illposed. To address this, we augment the learning objective with physics-based constraints–in the spirit of physics-informed neural networks (PINNs).^26–28^ Specifically, we regularize the reconstruction using the Helfrich-Canham-Evans bending energy,^29–31^ which describes the energetic cost of bending a membrane represented as a thin elastic sheet.^32,33^ By encouraging the reconstructed membrane to satisfy this physical model, the learned representation recovers geometrically smooth and physically plausible membrane surfaces, improving the accuracy of estimated membrane curvatures.

This study presents a physics-informed framework for reconstructing membrane geometry and its curvature distribution from volumetric imaging data. We combine INRs with Helfrich energy-based regularization to recover membrane geometries that are consistent with both experimental observations and membrane mechanics. The combination of INRs with Helfrich energy was previously explored by Rohanizadegan et al.,^20^ who used membrane mechanics to predict equilibrium vesicle shapes under prescribed physical parameters. Our study extends this approach to image-based membrane reconstruction, where the geometry is determined primarily by experimental observations, while membrane mechanics serves as a physical regularizer to improve curvature accuracy. A similar strategy of combining image-derived constraints with membrane mechanics has been employed in Helfrich Monte Carlo Flexible Fitting (HMFF),^34^ which reconstructs membrane surfaces using triangulated meshes. Unlike this mesh-based discrete approach, our framework employs a continuous INR, allowing geometric evaluation at arbitrary spatial resolutions and accommodating complex membrane topologies without prescribing mesh connectivity. The capabilities of our framework in comparison with existing methods for membrane segmentation, curvature estimation, simulation, and reconstruction are summarized in Table 1. Our work thus extends Helfrich-regularized INRs to image-based membrane reconstruction, enabling continuous representation of membrane geometry and accurate estimation of its curvature distribution.

**Table 1:** Comparison of representative methods for membrane segmentation, curvature estimation, membrane simulation, and reconstruction. The table summarizes the principal capabilities of each methodological category. ✓ indicates a supported or central capability; 1:, indicates partial, indirect, or implementation-dependent support; and – indicates that the feature is not a primary capability of the representative methods.

| **Methodological category** | **Membrane segmentation** | **Curvature estimation** | **Membrane simulation** | **Physics-informed fitting** | **Neural membrane mechanics** | **Neural image reconstruction** | **Physics-informed neural membrane reconstruction** |
| *Examples* | MemBrain-seg<sup>16</sup><br>TARDIS-em<sup>17</sup> | PyCurv<sup>21</sup><br>SurfORA<sup>37</sup><br>libigl<sup>38</sup> | Mem3DG<sup>39</sup><br>FreeDTS<sup>40</sup> | HMFF<sup>34</sup> | Rohanizadegan et al. (2024)<sup>20</sup> | vEMINR<sup>41</sup><br>ICE-TIDE<sup>42</sup><br>cryoDRGN<sup>43</sup> | This work |
|---|---|---|---|---|---|---|---|
| Image-based membrane identification | ✓ | – | – | △ | – | △ | △ |
| Geometry reconstruction from observations | △ | – | – | ✓ | – | △ | ✓ |
| Mean and Gaussian curvature estimation | – | ✓ | ✓ | ✓ | ✓ | – | ✓ |
| Continuous implicit surface representation | – | – | – | – | ✓ | △ | ✓ |
| Curvature via automatic differentiation | – | – | – | – | ✓ | – | ✓ |
| Helfrich-based membrane mechanics | – | – | ✓ | ✓ | ✓ | – | ✓ |
| Physics-constrained inverse reconstruction | – | – | – | ✓ | – | – | ✓ |
| Joint geometry–curvature optimization | – | – | △ | ✓ | ✓ | – | ✓ |
| Constrained by image-derived observations | ✓ | – | – | ✓ | – | ✓ | ✓ |
| Geometric representation | voxel | mesh | mesh | mesh | INR | INR | INR |

We evaluated the proposed framework through synthetic benchmarks and applications to experimental electron microscopy data. Using synthetic image data generated from mechanically minimized membrane meshes, we validated the accuracy of the reconstructed geometry and its differential properties, including surface normals and curvature. We further benchmarked the robustness of our framework against common three-dimensional electron microscopy (EM) artifacts including low signal-to-noise, anisotropic voxel resolution, and the missing wedge effect, showing that the inclusion of physics-based regularization improves the accuracy of the extracted geometries. These results establish a general framework for learning an implicit neural representation of membrane ultrastructure from images while supporting robust quantitative analysis of membrane ultrastructure and its functional consequences for cells.

As a practical application, we applied our framework to experimental electron microscopy datasets of mitochondria and the Golgi apparatus. The resulting three-dimensional segmentations of these organellar geometries were smooth across a wide range of membrane curvatures, illustrating the ability of our framework to learn the complex topology of these organelles without prior information and with little manual intervention required. We further applied our framework to endoplasmic reticulum–plasma membrane (ER–PM) contact sites to characterize local membrane geometries associated with intermembrane tethering, illustrating its potential for investigating biologically relevant membrane deformations. We also applied our framework to measure the distribution of membrane curvature at clathrin-mediated endocytosis sites. The extracted curvature profiles were azimuthally heterogeneous, which corroborated predictions from prior computational models showing a non-axisymmetric mechanical collapse of the membrane neck during clathrin-mediated endocytosis. Such coke-can-like buckling is thought to reduce the energy barrier for constricting the neck prior to vesicle hemifusion.^35,36^

## C Model formulation

The goal of our framework is to reconstruct membrane geometry from volumetric imaging data in a form that enables accurate estimation of differential geometric quantities, particularly membrane curvature (Fig. 1B). This objective can be distilled into two key requirements: (1) the reconstructed membrane geometry should be consistent with the volumetric image data, and (2) its curvature distribution should be consistent with a membrane mechanics model. In addition, the framework should (3) accommodate topologically complex membrane geometries without manual intervention, (4) remain robust to imaging noise and other experimental artifacts often encountered in volume electron microscopy datasets, and (5) enable direct computation of differential geometric quantities, such as curvature.

To satisfy these requirements, we formulate membrane reconstruction as an optimization problem over the membrane geometry. We represent the membrane using an implicit neural representation (INR).^24,25^ An INR represents a three-dimensional object as a continuous function parameterized by a multilayer perceptron (MLP). Because the network is differentiable, curvature and its derivatives can be evaluated directly and used in the loss functions. An additional advantage is that it naturally accommodates topological flexibility, satisfying our third objective. More specifically, we define a continuous scalar field which approaches values of −1 and 1 on either side of the membrane and whose zero level set defines the membrane surface (Fig. 1B).^20,44–46^ The formulation of the implicit neural representation is described in § G.1.

The membrane geometry, represented by the INR, is optimized by minimizing the following loss function:

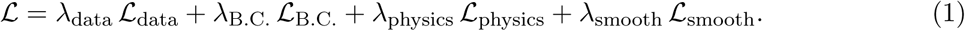

The first two terms, the data loss (*L*_data_) and boundary loss (*L*_B.C._), enforce consistency between the reconstructed membrane geometry and the membrane structure inferred from the volumetric image data, satisfying the first objective. The remaining terms, the physics loss (*L*_physics_) and smoothing loss (*L*_smooth_), regularize the reconstruction so that its curvature distribution is consistent with the membrane mechanics model, satisfying the second objective. The coefficients *λ*_data_, *λ*_B.C._, *λ*_physics_, and *λ*_smooth_ control the relative contributions of each loss term. Minimizing the total loss yields a reconstruction that balances agreement with the image data and physics-based regularization, rendering the approach robust to imaging noise and experimental artifacts, satisfying our fourth objective. The formulations of the individual loss functions and the optimization strategy are described in § G.2 and § G.3, respectively.

After optimization, the membrane surface is reconstructed from the optimized phase field, and its geometric properties, including the mean and Gaussian curvatures, are computed. Because the membrane is represented as an INR, these differential geometric quantities can be computed directly from the reconstructed field, satisfying the fifth objective. The postprocessing procedure is described in § G.4.

## D Results

### D.1 PINNs recover curvature-resolved membrane geometry from synthetic bench-marks

To quantitatively validate the proposed framework, we constructed synthetic membrane bench-marks that have a known geometric ground truth. Three representative membrane morphologies—a closed biconcave geometry, an open Ω-shaped budding geometry, and an open asymmetric protrusion geometry—were generated (Fig. 2A). The first two geometries were generated using Mem3DG,^39,47^ while the third was generated using FreeDTS,^40^ both of which are three-dimensional membrane simulators based on the Helfrich–Canham–Evans membrane model.^29–31^ These three geometries represent distinct mechanisms of membrane deformation: the biconcave geometry arises from zero spontaneous curvature under a prescribed enclosed volume and surface area, the Ω-shaped bud is generated by a heterogeneous distribution of spontaneous curvature, and the asymmetric protrusion is induced by the clustering of curvature-generating proteins. These Helfrich-based geometries provide suitable benchmarks for our PINN framework, which is designed to reconstruct cellular membranes whose mechanical behavior can be approximated by the same energy formulation. Notably, the FreeDTS configuration incorporates thermal fluctuations and protein-induced deformations, providing a benchmark that does not need to satisfy the Helfrich energy minimization assumption and allowing us to evaluate reconstruction accuracy under physically plausible deviations from the assumed physical prior. We converted the synthetic mesh geometries into volumetric data and applied synthetic noise to mimic realistic electron tomography images (Fig. 2B, top). We then extracted point signals from the images to define the collocation points used to compute the loss functions (Fig. 2B, remaining panels).

**Figure 2:**
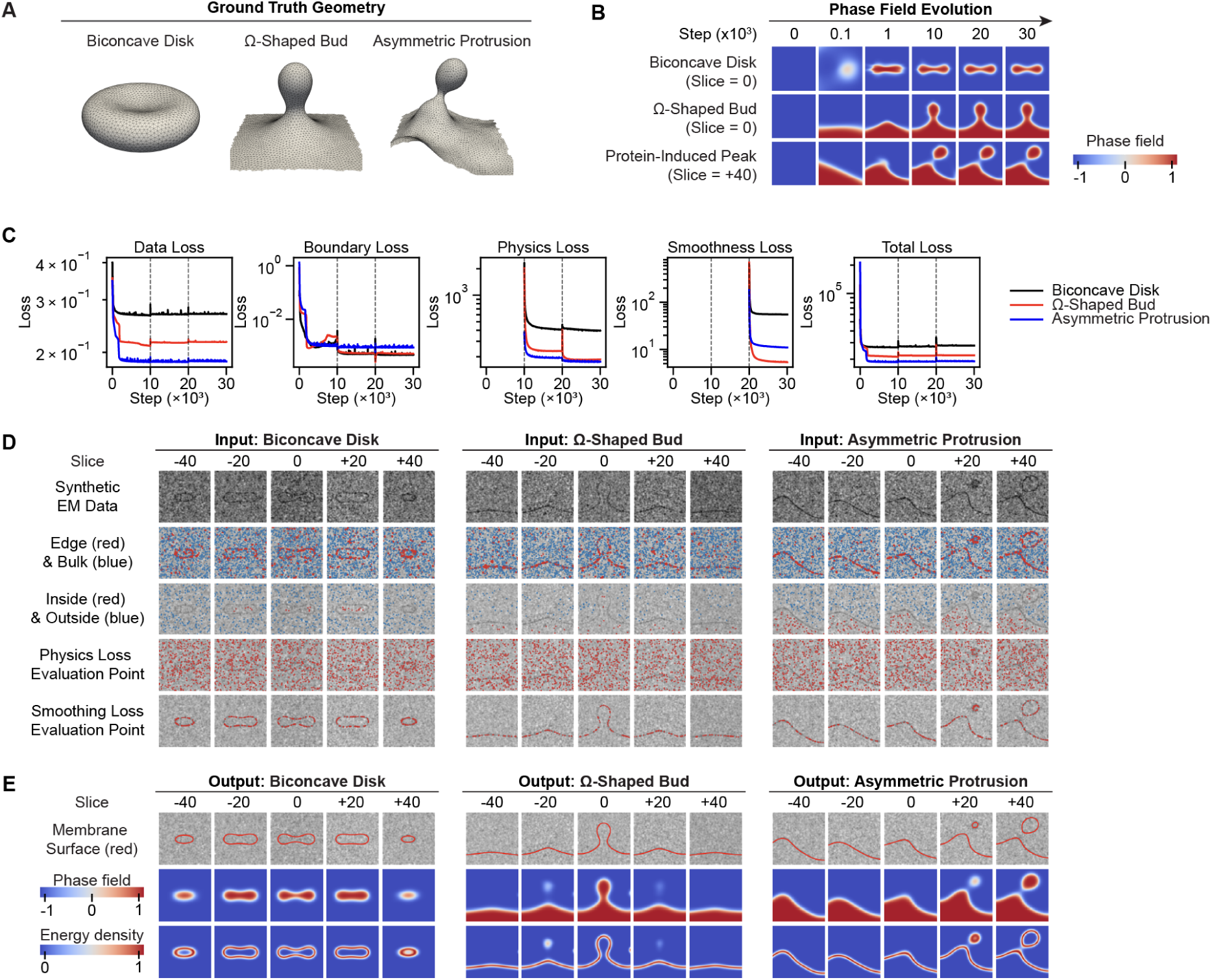
Optimization on a synthetic membrane dataset converges to a stable solution that accurately reproduces the membrane geometry. (A) Ground truth membrane shapes generated using three-dimensional membrane mechanics simulator, Mem3DG and FreeDTS. (B) Evolution of the phase field across optimization steps. (C) Evolution of the loss functions over optimization steps. Physics and smoothing losses are introduced at steps 10 × 10^3^ and 20 × 10^3^, respectively. (D) Input voxel data and sampled point signals utilized in the evaluation of model loss by image slice. (E) Final reconstruction shown slice by slice. Top: reconstructed membrane surface overlaid on synthetic EM data. Middle: phase field *φ*. Bottom: energy density *eE*.

We first evaluated the stability and convergence of the proposed optimization framework. Initial efforts to simultaneously optimize all four loss functions led to poor and unstable model convergence, likely because the physics and smoothing losses depend on high-order derivatives of the phase field, which are poorly behaved when the field is far from the target solution. We therefore adopted a three-stage optimization strategy (Fig. 1C). In the first stage, only the data and boundary losses were active, aligning the phase field with the target membrane geometry. The physics loss was introduced in the second stage to stabilize the differential properties of the phase field, followed by the curvature-smoothing loss in the third stage to refine the reconstructed curvature. With this staged optimization strategy, all loss components converged stably throughout training, with no observable instability following the introduction of the additional objectives (Fig. 2C). As optimization progressed, the phase field evolved from a uniform scalar field into a smooth implicit representation (Fig. 2D) that recovered the target membrane morphology (Fig. 2E).

We next evaluated the accuracy of the reconstructed membrane geometry and its differential geometric properties after each optimization stage. Following the first stage, the reconstructed membrane geometry approximated the ground truth geometry (Fig. 3A). This agreement was quantified using the surface Dice score, which measures the overlap between reconstructed and ground truth surfaces (Fig. 3C). Surface Dice remained near 1.0 at every stage, improving modestly once the physics and smoothing losses were added. We further evaluated the accuracy of reconstructed surface normal angles, mean curvatures, and Gaussian curvatures by sampling points on the re-constructed membrane surface and comparing their geometric quantities with those of the nearest neighbors on the ground truth surface (Fig. 3A). While the surface normal angle showed close agreement with the ground truth after the first stage, mean and Gaussian curvatures exhibited poor correlations (Fig. 3B and C). Sequential incorporation of the physics and smoothing losses during the second and third optimization stages improved the agreement of both mean and Gaussian curvatures with the ground truth (Fig. 3B and C). By the final stage, all geometric quantities exhibited strong correlations with the ground truth (*r >* 0.95). To confirm that these improvements were not simply due to additional optimization iterations, we performed controlled ablation experiments with matched optimization budgets (Fig. S3), demonstrating that prolonged optimization alone did not improve curvature accuracy. These results demonstrate that the data and boundary losses enable the reconstruction of the membrane’s general location in space, while the physics and smoothing terms enable the accurate reconstruction of higher-order differential geometric quantities, including mean and Gaussian curvature. To further investigate the role of physics regularization in achieving this improvement, we performed an ablation study in which the physics loss was omitted from Stage 2 (*λ*_physics_ = 0), while retaining the curvature-smoothing loss in Stage 3. Without physics regularization, the subsequent curvature-smoothing optimization became unstable for some geometries, resulting in numerical failure or substantial deformation of the reconstructed membrane surface (Fig. S2A-B). The higher-order spatial derivatives of the phase field were larger and less stable without the physics loss (Fig. S2C-D). These results indicate that Stage 2 serves not only to improve curvature estimation directly, but also to stabilize the higher-order differential properties of the phase field, providing a more well-conditioned representation for subsequent curvature optimization in Stage 3.

**Figure 3:**
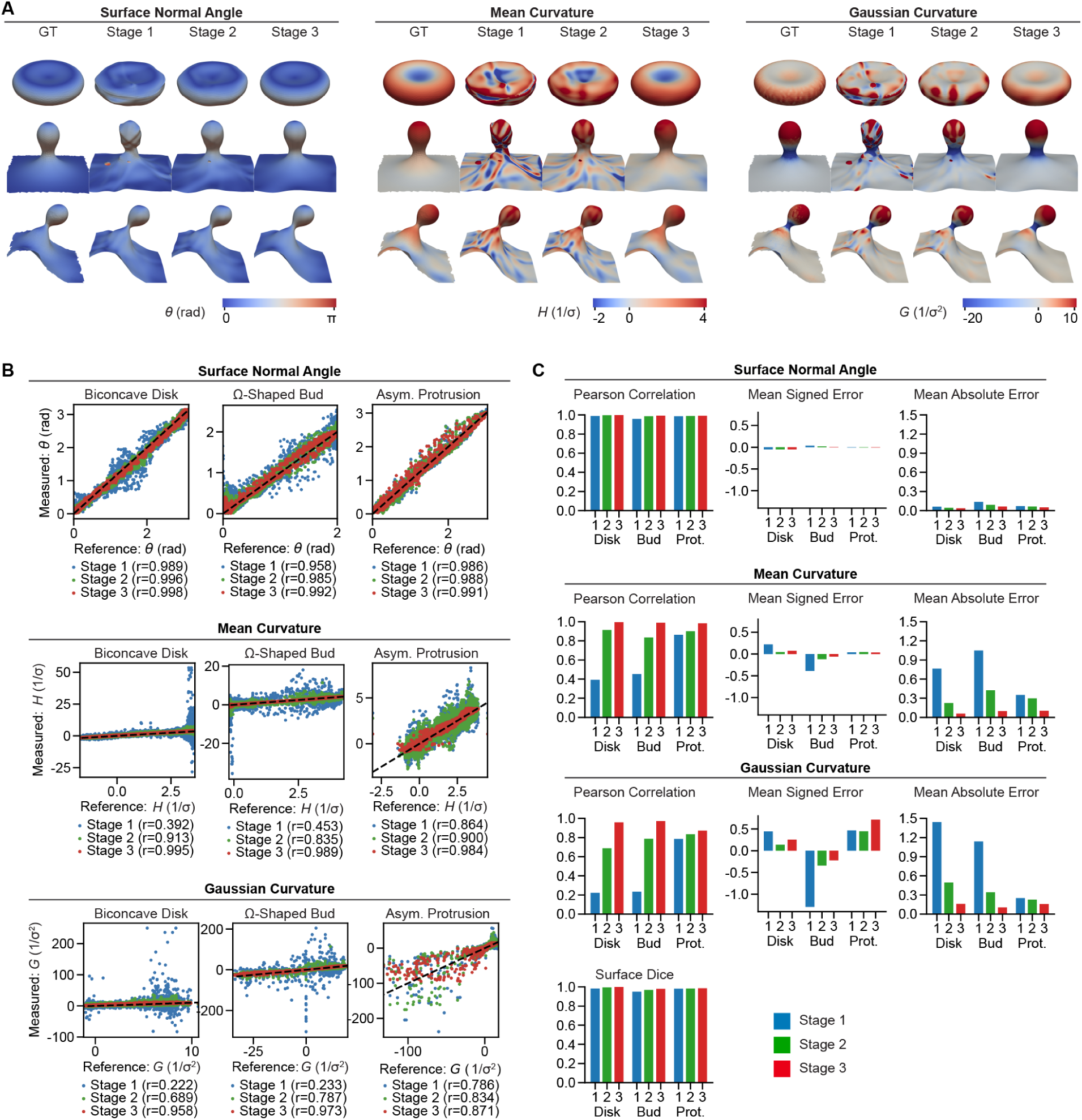
Physics-based regularization enables accurate recovery of membrane curvature beyond surface localization. (A) Three-dimensional reconstruction of membrane geometry and associated geometric quantities. Surface color maps indicate the surface normal angle, mean curvature, and Gaussian curvature. Reconstructions after each optimization stage are shown alongside the corresponding ground truth (GT). (B) Correlation analysis between ground truth and reconstructed geometric quantities. The dashed line indicates the identity line (slope = 1), and Pearson correlation coefficients (*r*) are reported for each case. (C) Performance metrics at each optimization stage. Shown are dice score, surface normal correlation, mean curvature correlation, and Gaussian curvature correlation for the biconcave and bud synthetic datasets after each stage of the optimization.

Finally, using the biconcave disk geometry, we investigated the sensitivity of reconstruction accuracy and stability to the loss weights (Fig. S4). The data and physics loss weights exhibited a trade-off between data fidelity and physical regularization. Decreasing the data loss weight or increasing the physics loss weight placed greater emphasis on satisfying the physical constraints, resulting in overly rounded geometries that failed to capture morphological details. Conversely, increasing the data loss weight or decreasing the physics loss weight increased sensitivity to noise, frequently leading to reconstruction instability. Increasing the boundary loss weight had relatively little effect on reconstruction accuracy, whereas decreasing it weakened the phase-field sign constraints and could result in reconstruction failure. The smoothing loss weight influenced both curvature accuracy and reconstruction stability: insufficient smoothing degraded curvature estimation, whereas excessive smoothing led to reconstruction failure. These results highlight the importance of balancing the individual loss contributions to achieve accurate and stable membrane reconstruction.

Together, these results demonstrate that the proposed PINN framework can recover membrane geometry with curvature-level fidelity through staged optimization and appropriately balanced loss contributions.

### D.2 PINNs enable robust curvature recovery from noisy volumetric data

Having established the effectiveness of PINN for geometric reconstruction, we next asked whether the PINN framework is robust to imaging noise. In practice, volume imaging datasets produced via approaches such as three-dimensional electron microscopy have a poor signal-to-noise ratio which can hinder quantitative analysis workflows. We assessed the robustness of the PINN framework to imaging noise by evaluating reconstruction performance across a series of synthetic datasets with increasing levels of noise (Fig. 4A).

**Figure 4:**
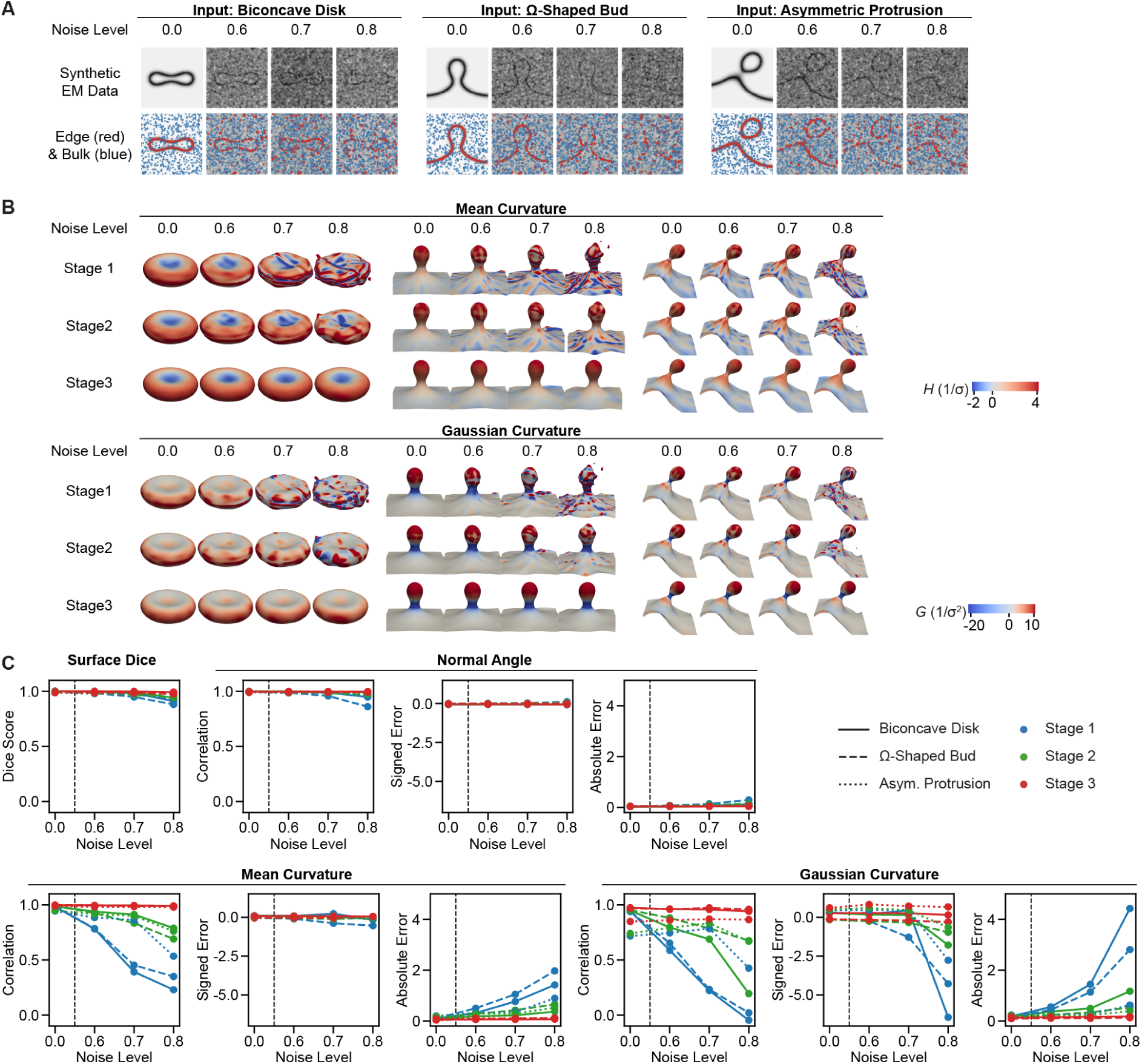
Curvature calculation remains robust under imaging perturbations through physics-based regularization. (A) Synthetic EM images and extracted edge/bulk signals with increasing noise. (B) Three-dimensional reconstruction of membrane shape and curvature distribution at each optimization stage and across different noise levels. (C) Quantitative comparison with the ground truth. Surface agreement was evaluated using the surface Dice score, while surface normal angle, mean curvature, and Gaussian curvature were assessed using Pearson correlation coefficients, mean signed error, and mean absolute error.

Specifically, for each image in the series a prescribed fraction (referred to as the “noise level”) of membrane voxels was randomly removed, while a fixed fraction of background voxels was randomly reassigned as membrane prior to Gaussian filtering and normalization. Increasing the noise level from 0.0 to 0.8 degraded the clarity of membrane surface boundaries in the synthetic images (Fig. 4A). Across all noise conditions, the edge/bulk signals were derived using the standard unmodified thresholding procedure, resulting in reduced signal accuracy as noise increased. In contrast, the inside/outside signals were kept fixed, reflecting our experimental workflow in which a small number of confidently labeled points are manually selected based on visual inspection and are therefore less sensitive to image degradation. A separate sensitivity analysis (Fig. S5) showed that reconstruction can remain accurate with relatively few inside/outside points, provided that these points are correctly labeled and sufficiently distributed, supporting the validity of this approach under noisy imaging conditions. We applied the PINN framework to each dataset and evaluated reconstruction accuracy using surface Dice to quantify morphological agreement with the ground truth. We also assessed the accuracy of surface normal orientations, mean curvatures, and Gaussian curvatures using Pearson correlation coefficients, normalized signed errors, and normalized absolute errors relative to the ground truth.

Our result showed that the surface Dice remained high across all noise levels and optimization stages (Fig. 4C), indicating that the overall membrane morphology was preserved even when the input images were very degraded. Surface normal orientation also showed strong agreement with the ground truth across all conditions. In contrast, in spite of the apparent morphological accuracy, the correlation of reconstructed mean and Gaussian curvatures with the ground truth was reduced with increasing noise level (Fig. 4B and C). Incorporation of the physics and smoothing losses in stages 2 and 3 improved the accuracy of the reconstructed quantities. By stage 3, both mean and Gaussian curvature showed strong agreement with the ground truth even under high noise levels, leading us to conclude that the PINN framework recovers curvature-level geometric information from noisy images.

In addition to image noise, three-dimensional electron microscopy approaches are also subject to other imaging artifacts such as voxel anisotropy and the missing wedge effect. To further assess the ability of PINN to recover curvature under realistic imaging artifacts, we extended the analysis to two additional data loss conditions commonly observed in experimental datasets. First, we introduced voxel anisotropy by increasing the voxel size along the *z* direction relative to the *x* and *y* directions by up to a factor of 8 (referred to as “anisotropy”, Fig. S6). Increasing anisotropy effectively reduced the resolution along the *z* axis, leading to step-like artifacts in the reconstructed surface after stage 1. These artifacts were reduced by physics-based regularization in stage 2, and the smoothing loss in stage 3, resulting in improved curvature estimation. Second, we simulated the missing wedge effect by restricting the tilt angle range from 90° to 60° (Fig. S7). As the tilt range decreased, large regions of the membrane had missing intensity in the simulated images, leading to incomplete input data. This posed a larger challenge for reconstruction, resulting in global morphological errors. While the physics and smoothing constraints reduced the roughness of reconstructed surfaces, they were insufficient to recover the missing global structure when entire regions of the membrane were absent. These results suggest that the PINN framework is robust to local degradation in data quality but remains limited by large-scale missing information.

Taken together, these results demonstrate that the PINN framework enables accurate recovery of curvature distributions under a range of realistic data imperfections. When input corruption introduces local distortions, such as noise or anisotropic resolution, physics and smoothing constraints restore reliable estimates of mean and Gaussian curvature by enforcing global geometric consistency. However, when large portions of the membrane are not present, as in the missing wedge condition, curvature cannot be accurately recovered due to the lack of global structural information. These results establish PINNs as a robust framework for curvature-resolved membrane reconstruction from imperfect volumetric data.

### D.3 PINNs reconstruct topologically complex membrane geometries from experimental volumetric data

To evaluate whether PINNs generalize beyond the synthetic benchmark geometries presented above, we next applied the framework to experimentally acquired electron microscopy data containing organelle membrane structures with complex topology. In contrast to the simple closed and open geometries used in the synthetic studies, cellular organelles have sophisticated morphologies with multiple enclosed compartments. Such topological complexity poses a major challenge for conventional mesh-based reconstruction pipelines, often requiring topology handling, mesh repair, or manual post-processing. In practice, these difficulties have led many studies to rely on labor-intensive manual segmentation of complex membrane structures. By representing membrane geometry as a continuous implicit field, PINNs can accommodate these complex topologies without explicit topology tracking or mesh surgery.

We applied PINNs to publicly available volumetric electron microscopy datasets of the Golgi apparatus in *Chlamydomonas reinhardtii*^48^ and neuronal mitochondria in the mouse cerebellum^49^ imaged by cryo-electron tomography and serial-section electron tomography, respectively. These organelles were selected as representative examples of membranes with complex architecture and topology; the Golgi apparatus consists of interconnected cisternal membranes with fenestrated sheet-like structures, whereas mitochondria exhibit nested double-membrane architectures with highly folded cristae. We extracted edge/bulk signals using a thresholding method for all slices, and inside/outside signals by manual inspection of several slices (Fig. 5A and B). Using these signals as input, we applied the three-stage optimization procedure, which reconstructed smooth three-dimensional membrane surfaces (Fig. 5C–F). Notably, after point-signal extraction, the remaining reconstruction process was automated, and explicit topology tracking or mesh repair was not needed.

**Figure 5:**
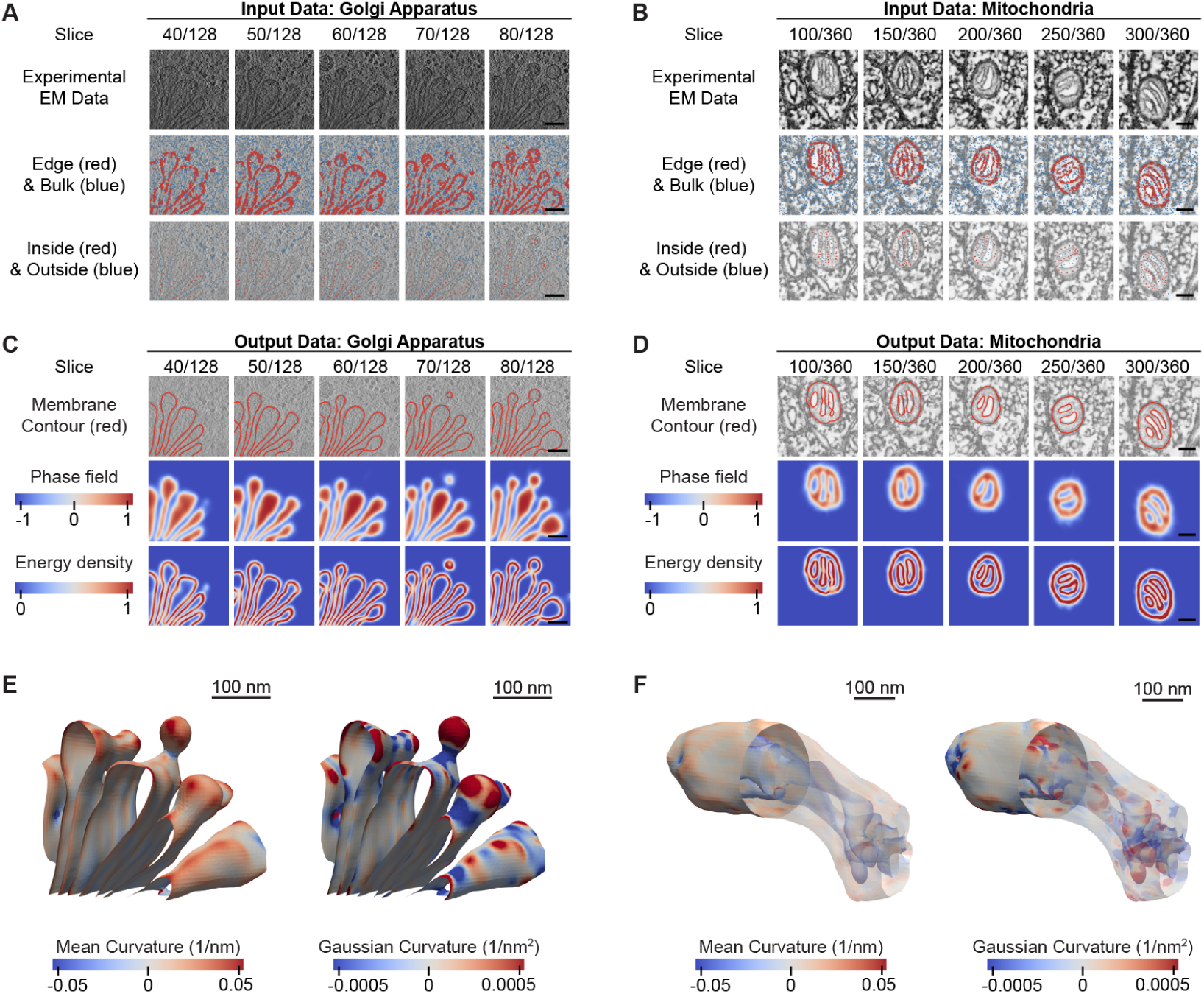
PINNs handle topologically complex organelle membrane structures from three-dimensional electron microscopy datasets of cells. Left: cryo-electron tomogram of *Chlamydomonas reinhardtii* after focused ion beam milling,^48^ showing the Golgi apparatus. Right: serial-section electron tomogram of mouse cerebellum after high-pressure freezing and freeze substitution,^49^ showing a mitochondrion. (A, B) Input EM images with extracted point-based signals, shown slice by slice. (C, D) Corresponding PINN reconstructions, shown slice by slice. Top: membrane contour (red) overlaid on the EM data. Middle: phase field *φ*. Bottom: energy density *eE*. (E, F) Three-dimensional membrane reconstructions colored by mean and Gaussian curvature. Scale bars represent 100 nm in all panels.

As observed in the overlay between the reconstructed surfaces and the original images, the reconstructed membranes matched the visible membrane structures in the input raw images (Fig. 5C and D). Three-dimensional visualization further confirmed that the reconstructed surfaces preserved topologically complex features, including stacked cisternal membranes and fenestrations in the Golgi apparatus, as well as the nested membrane architecture and branch–fusion structures in the mitochondrion (Fig. 5E and F). The resulting continuous and differentiable representation enabled the computation of mean and Gaussian curvatures across these topologically complex membrane geometries through automatic differentiations (Fig. 5E and F).

To quantitatively assess the geometric accuracy of the reconstructed membranes, we compared the PINN reconstructions with publicly available reference segmentation data (Fig. S10 and Table S1). For the Golgi apparatus, we used a voxel-based reference segmentation generated using MemBrain-seg^16^ and available through the CZII Data Portal,^48,50^ whereas for the mitochondrion, we used manual contour annotations provided by Mendelsohn et al.,^49^ defined on individual image slices. The Golgi reconstruction showed good agreement with the reference segmentation, with surface Dice scores of 0.815 and 0.966 at distance tolerances of 5 and 10 nm, respectively. Here, the distance tolerance is the threshold which defines the maximum separation between surfaces for them to be considered matching. The average surface distance (the cumulative mean of one-sided nearest neighbor distances in both directions) was 3.02 nm, comparable to the segmentation voxel spacing of 3.136 nm. The maximum and 95th percentile Hausdorff distances were 28.8 and 10.61 nm, respectively, reflecting localized discrepancies near the z-stack boundaries (Fig. S10B), where membrane edge signals became ambiguous. For the mitochondrion, the surface Dice scores were lower, reaching 0.612 and 0.840 at tolerances of 5 and 10 nm, respectively, with an average surface distance of 9.10 nm. These larger discrepancies may be attributed to two factors. First, the coarser image resolution and two regions of missing data from tissue lost during ultramicrotomy which introduce uncertainties in both the PINN reconstruction and manual segmentation. Second, the PINN reconstruction treated the closely spaced outer and inner mitochondrial membranes as a single membrane surface through thresholding-based signal extraction, whereas the manual segmentation traced them separately, with the outer membrane used as the reference here. This difference resulted in a systematic positional offset of the mitochondrial outer boundary (Fig. S10D), contributing to the larger average surface distance and Hausdorff distances of 62.0 nm (maximum) and 20.4 nm (95th percentile). Overall, these quantitative comparisons support the geometric accuracy of the PINN reconstructions, while revealing localized discrepancies associated with image quality and membrane signal extraction. These results demonstrate that our PINN framework is applicable also to real experimental data with membrane structures with greater topological complexity. The topology-agnostic implicit representation enables automated reconstruction and differential geometric analysis of membrane architectures that are challenging to handle using conventional mesh-based approaches.

### D.4 PINNs identify putative ER–PM tethering sites through membrane curvature analysis

To demonstrate the biological applicability of curvature estimation using PINNs, we analyzed endoplasmic reticulum–plasma membrane (ER–PM) contact sites in budding yeast (*Saccharomyces cerevisiae*). Previous studies have shown that ER–PM tethering proteins, particularly tricalbins, induce highly curved membrane protrusions on the cortical ER facing the plasma membrane.^51^ These localized curvature features provide a potential morphological signature of ER–PM tethering sites.

We applied our PINN framework to a cryo-electron tomogram of ER–PM contact sites (EMD-10378) from the same study.^51^ We first segmented the membrane using Membrainseg, based on which, we extracted edge/bulk and inside/outside point signals (Fig. 6A,B). We then reconstructed the membrane geometry using our three-step optimization (Fig. 6C,D). We then calculated the mean and Gaussian curvatures of the reconstructed surface (Fig. 6E,F).

**Figure 6:**
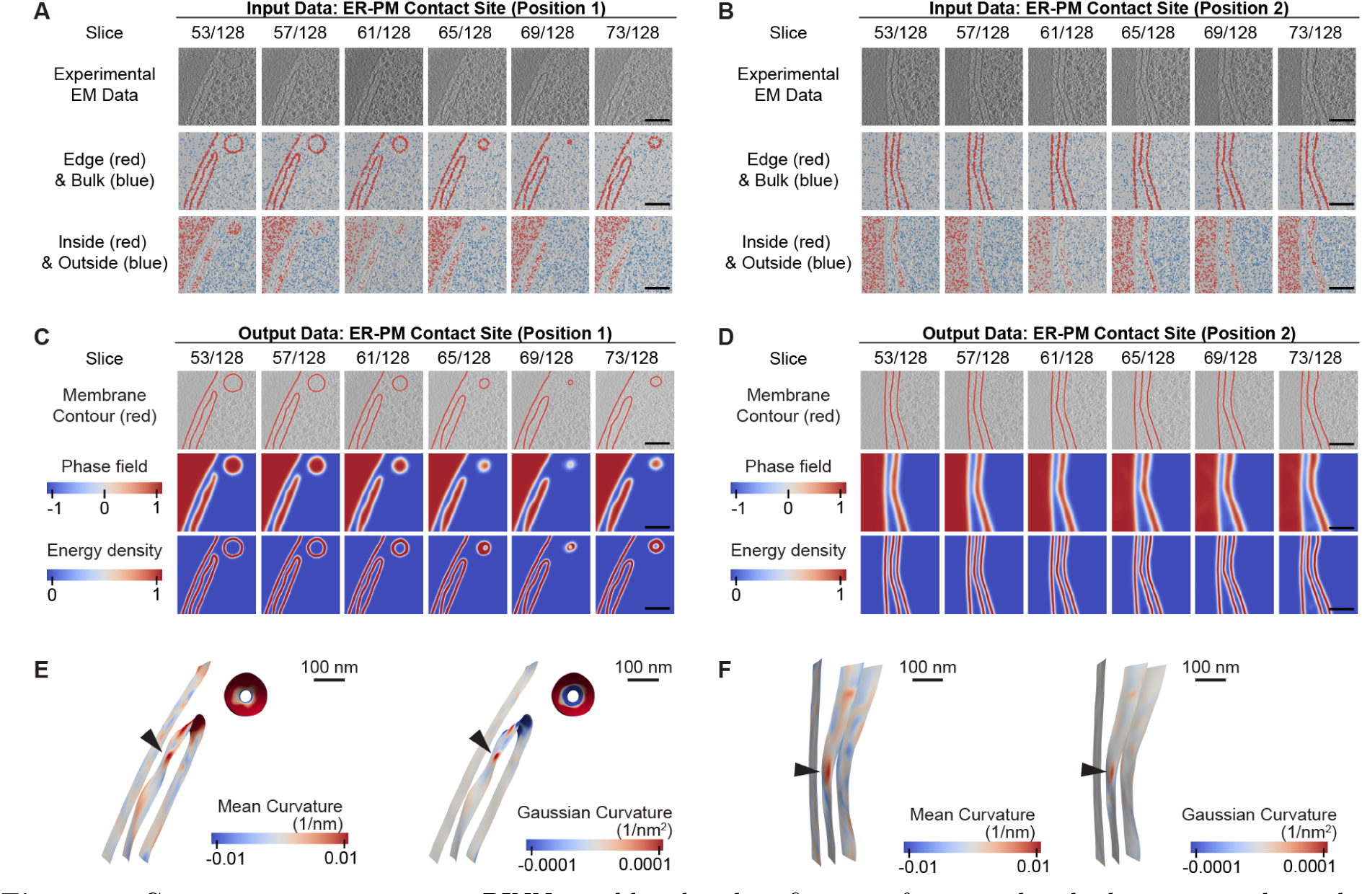
Curvature estimation using PINNs enables the identification of potential endoplasmic reticulum–plasma membrane (ER–PM) tethering sites. Data: Cryo-electron tomogram of budding yeast (*Saccharomyces cerevisiae*), prepared by cryo-focused ion beam milling, showing endoplasmic reticulum–plasma membrane (ER–PM) contact sites (EMD-10378).^51^ (A, B) Input EM images with extracted point-based signals, shown slice by slice. (C, D) Corresponding PINN reconstructions, shown slice by slice. Top: membrane contour (red) overlaid on the EM data. Middle: phase field *φ*. Bottom: energy density *eE*. (E, F) Three-dimensional membrane reconstructions colored by mean and Gaussian curvature. The black arrow indicates a putative ER–PM tethering site. Scale bars represent 100 nm in all panels.

The reconstructed curvature maps revealed spatially heterogeneous curvature distributions. Notably, localized regions of high curvature were observed on the cortical ER membrane facing the plasma membrane, as indicated by the black arrows in Fig. 6E and F. These features are consistent with the high-curvature membrane protrusions previously associated with tricalbin-mediated ER–PM tethering.^51^ Importantly, this result suggests that PINNs can retain localized high-curvature features despite physics and smoothing regularization, provided that these features are sufficiently captured by the image-derived data constraints. This example further illustrates how continuous curvature analysis using PINNs can reveal morphological signatures associated with molecular mechanisms of membrane deformation.

### D.5 PINNs reveal non-axisymmetric 3D curvature heterogeneity at endocytic pits

To demonstrate that PINNs enable differential geometric analysis in a biologically relevant context, we applied the framework to three-dimensional EM datasets of endocytic pits in cells. Endocytic pits are transient membrane invaginations formed during clathrin-mediated endocytosis, where progressive membrane deformation leads to the formation of a narrow neck prior to vesicle scission.^53^ Mechanical simulations of membrane remodeling have predicted that anisotropic curvature distributions can emerge at the neck region as energetically favored equilibrium configurations.^35,36^ Although previous experimental studies have characterized endocytic pit morphology using profile analysis of electron tomography data,^6^ these analyses primarily reduced membrane geometry to rotationally symmetric profiles, leaving how curvature varies across the full three-dimensional neck geometry largely unknown. PINNs provide a continuous and differentiable representation of three-dimensional membrane geometry, enabling quantitative analysis of curvature distributions across different endocytic pit stages.

We applied PINNs to a publicly available dataset of COS-7 cells prepared by high-pressure freezing and freeze-substitution, and imaged by FIB-SEM.^14,52^ From this dataset, we selected five endocytic sites for analysis. PINNs successfully reconstructed smooth three-dimensional membrane surfaces from all sites (Fig. 7A and B), enabling direct computation of mean and Gaussian curvatures over the reconstructed geometries (Fig. 7D and E). As expected for a budding geometry, mean curvature increased toward the bud tip and Gaussian curvature was negative at the neck. However, unlike axisymmetric idealizations, the curvature maps varied along the azimuthal direction (i.e., around the neck).

**Figure 7:**
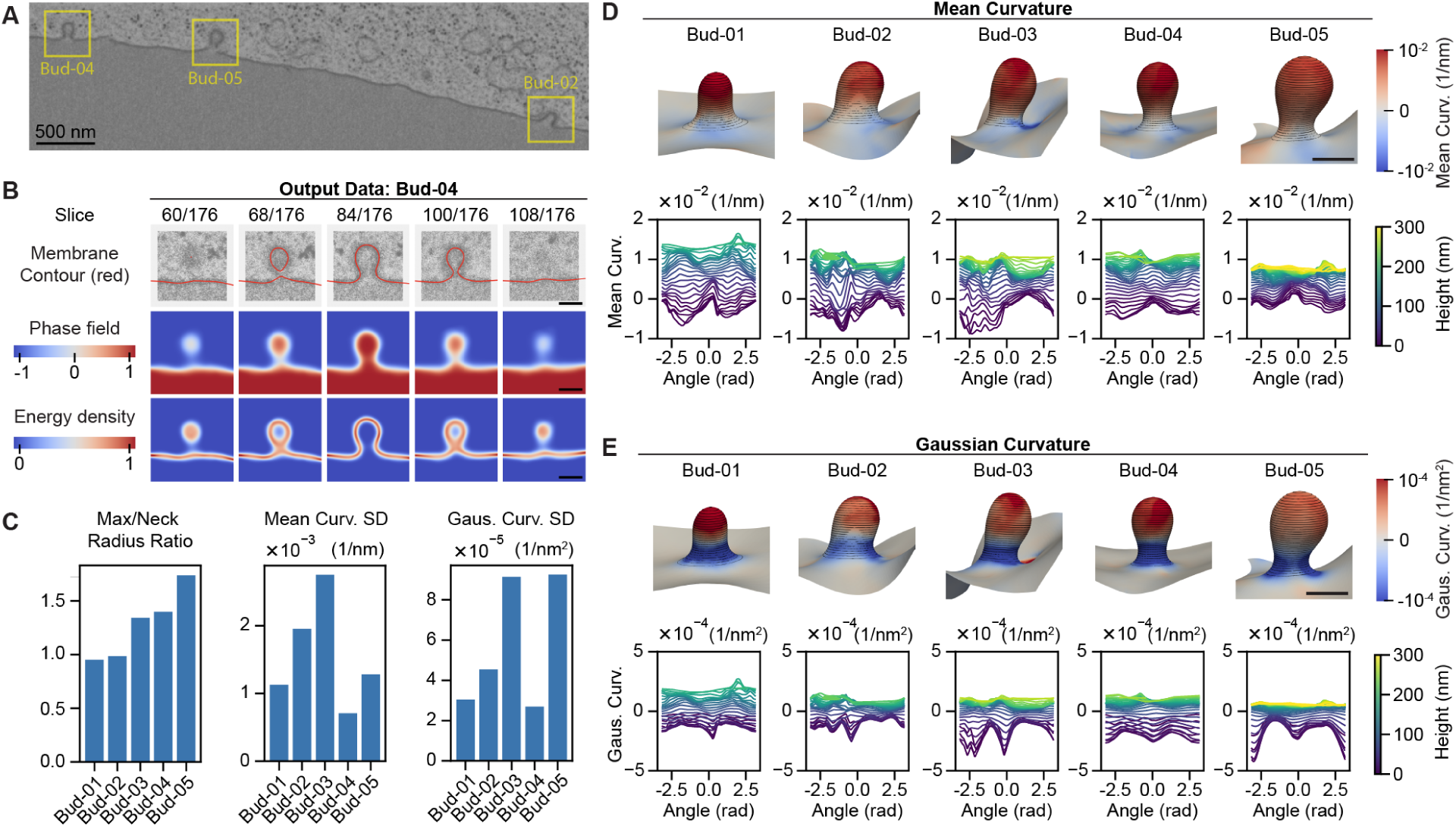
PINNs reveal three-dimensional variation in membrane curvature at the necks of endocytic pits. (A) Example FIB-SEM image of COS-7 cells after high-pressure freezing and freeze substitution.^14,52^ Manually identified endocytic pits are outlined in yellow. (B) PINN reconstruction of Bud-04, shown slice by slice. (C) Maximum-to-neck radius ratio and standard deviations of mean and Gaussian curvature for five endocytic pits. (D) Mean curvature distribution and its quantification along each contour ring. (E) Gaussian curvature distribution and its quantification along each contour ring. Scale bars represent 500 nm in panel A and 100 nm in panel B, D, and E.

To quantitatively characterize these curvature variations, we sampled contours at different heights relative to the pit base and traced the profiles of mean and Gaussian curvatures around each ring (Fig. 7D and E). As the contours approached the neck region, both mean and Gaussian curvatures exhibited increasing curvature azimuthal heterogeneity, indicating a progressive anisotropy in neck curvature. To assess the sensitivity of these observations to curvature regularization, we repeated the reconstructions with different smoothing loss weights, *λ*_smooth_ = 1, 5, 15 (Fig. S11). The spatial patterns of curvature heterogeneity remained consistent across the tested weights, suggesting that the observed anisotropy is not an artifact of the specific smoothing weight used in the reference reconstruction.

We further examined the relationship between curvature variation and endocytic progression (Fig. 7C). We ordered the pits according to the ratio of the maximum fitted contour-ring radius to the neck radius, a proxy for the progress of endocytosis.^6^ We first reported how the mean curvature varies at the neck of pits at different stages of endocytic progression. The necks of these endocytic pits showed variability in mean curvature, especially at intermediate stages of endocytic progression (Figure 7C and D). We next quantified the variance in Gaussian curvature at the neck region, as Gaussian curvature is particularly sensitive to anisotropic saddle-like deformation of the neck. Gaussian curvature of endocytic necks varied azimuthally, with the largest variance in neck Gaussian curvature observed in pits at intermediate and late stages (Figure 7C and E).

These results demonstrate that PINNs enable differential geometric analysis directly from experimentally acquired volumetric electron microscopy data, providing access to geometric quantities beyond surface morphology alone. In particular, the analysis revealed anisotropic curvature at the necks of intermediate- and late-stage endocytic pits, consistent with membrane deformation be-coming increasingly spatially heterogeneous during pit maturation. By reconstructing membrane geometry as a continuous and differentiable field, PINNs enable quantitative characterization of local curvature distributions, azimuthal heterogeneity, and stage-dependent geometric variations that are difficult to resolve using conventional voxel-based representations. More broadly, this frame-work provides a general approach for linking volumetric electron microscopy data with differential geometric and mechanical analyses of dynamic membrane remodeling processes.

## E Discussion

Through systematic validation on synthetic datasets, we demonstrated that the proposed frame-work robustly recovers curvature distributions even under lossy input conditions. Application to experimental datasets further showed that the method generalizes to complex cellular structures, including the Golgi apparatus and mitochondria, without requiring prior specification of membrane topology. We also applied our method to the analysis of endocytic pits and performed three-dimensional curvature analysis, revealing non-axisymmetric curvature distributions at the neck region. These results highlight the potential of our approach to uncover structural features that are difficult to resolve using conventional methods.

### E.1 Related work

We summarized the related work in Table 1 together with the capacity of each method.

Among existing methods, HMFF^34^ is most closely related to our PINN framework in its aim of reconstructing membrane geometries using both image data and physical constraint. The primary distinction lies in the representation of the membrane. HMFF uses an explicit triangulated mesh, whereas our framework represents the membrane as the zero level set of a continuous phase field parameterized by an INR. This distinction determines how geometric quantities are computed. In HMFF, curvature is computed from the local geometry of the mesh, while in our framework, surface normals and curvatures are evaluated directly from spatial derivatives of the learned phase field. The two methods also differ in how they incorporate imaging data and membrane mechanics into the fitting procedure. HMFF samples mesh configurations through Monte Carlo simulations guided by the Helfrich bending energy and an image-derived bias. Our PINN framework fits the phase field by minimizing a composite loss that combines data fidelity with Helfrich-based regularization. The choice of representation also affects how membrane topology is specified. HMFF starts from a mesh with prescribed connectivity, which can evolve when topology-changing moves are enabled. In our framework, membrane topology emerges from fitting the continuous phase field to image-derived edge/bulk and inside/outside signals, without the need to prescribe an initial topology.

Several prior studies have quantified membrane curvature from reconstructed membrane geometries in electron microscopy data. Methods such as PyCurv,^21^ surface morphometrics,^23^ SurfORA,^37^ and libigl^38^ estimate curvature from explicit triangular meshes. PyCurv and surface morphometrics employ a tensorvoting framework to estimate curvature over geodesic surface neighborhoods, whereas SurfORA and libigl use local surface fitting to estimate principal curvatures from neighboring mesh vertices. Mem3DG,^39^ a framework for simulating membrane mechanics, computes curvature directly from triangulated meshes using a discrete differential geometry formulation. Despite differences in implementation, these approaches rely on discrete local geometry derived from reconstructed surface meshes. To quantitatively compare these approaches, we evaluated curvature estimation accuracy using synthetic geometries with known ground truth under varying mesh resolutions and levels of vertex perturbation (Figs. S8 and S9). The results showed that mesh-based methods generally achieved accurate curvature estimates comparable to those of our PINN frame-work when high-quality meshes were available, although their accuracy deteriorated for coarse or strongly perturbed meshes. The results showed that mesh-based methods generally achieved accurate curvature estimates when high-quality meshes were available, although their accuracy decreased for coarse or strongly perturbed meshes. Our PINN framework achieved comparable curvature accuracy while employing a fundamentally different geometric representation, in which membrane geometry is described by a continuous differentiable implicit neural field and curvature is computed directly from its spatial derivatives. Thus, our framework provides a complementary approach to established mesh-based methods, enabling membrane reconstruction and curvature analysis directly from image-derived signals without requiring an intermediate mesh representation.

INRs have recently emerged as a powerful framework for cryo-electron microscopy and tomography reconstruction.^24,25^ To date, however, they have been applied mainly to the problem of recovering a three-dimensional density field from imaging data, rather than to model the geometry of the imaged structures themselves. In single-particle cryo-EM, foundational methods such as cryoDRGN^43^ and subsequent neural-field approaches^54–56^ have used continuous neural representations to reconstruct volumetric density while modeling structural heterogeneity and conformational variation. More recently, INR-based methods have been extended to cryo-electron tomography. ICE-TIDE^42^ jointly optimizes volumetric reconstruction together with deformation and alignment correction from tilt-series data, while quantEM^57^ employs INRs for missing wedge inpainting and joint tomographic alignment. Similar ideas have also been applied to volume electron microscopy, where vEMINR^41^ represents large-scale EM volumes as continuous neural fields, enabling compact volumetric representation together with interpolation and super-resolution. Importantly, these approaches focus on density reconstruction and imaging-related inverse problems. Our method instead uses an implicit neural representation as a membrane-specific phase field, where the learned continuous field defines membrane geometry and enables analytical computation of differential geometric quantities through spatial derivatives.

### E.2 Applicability to experimental datasets

In this study, we demonstrated the applicability of our method to experimental membrane data using four distinct examples: the Golgi apparatus, mitochondria, an ER-PM contact site, and endocytic pits (Fig. 5 – Fig. 7). These datasets were acquired using different electron microscopy modalities, namely *in situ* cryo-electron tomography,^48^ serial-section electron tomography,^49^ and focused ion beam scanning electron microscopy,^14,52^ respectively. Beyond differences in three-dimensional EM modality, the sample preparation across these studies also differed. The Golgi dataset was acquired from vitrified specimens without chemical staining,^48^ whereas the mitochondria and endocytic pit datasets were obtained from resin-embedded samples enhanced with heavy-metal contrast agents.^14,49,52^ Despite these differences in imaging modality, sample preparation, and image contrast, our method successfully reconstructed smooth membrane geometries with curvature-level resolution across all datasets. These results suggest that the proposed framework is broadly applicable to diverse experimental membrane imaging data obtained under a wide range of acquisition conditions.

Compared with existing membrane segmentation and reconstruction approaches, the advantages of our method can be viewed from two perspectives: the accuracy of higher-order geometric quantities and the reduced need for human intervention. Two of the experimental datasets used in this study, namely the Golgi apparatus and mitochondria, have previously been analyzed using alternative reconstruction approaches, enabling direct comparison on the same biological specimens. For the Golgi apparatus dataset,^48^ membrane masks segmented by a convolutional neural network (CNN)-based semantic segmentation model^58^ and MemBrain-seg^16^ are publicly available.^50^ The general CNN-based semantic segmentation model captures membrane localization on a slice-by-slice basis, but often exhibits limited continuity along the *z*-direction. This issue is substantially improved in MemBrain-seg, where the membrane masks appear more continuous across all spatial directions. While these segmentation results appear visually comparable to our reconstructions, their voxel-based representation inevitably introduces discretization artifacts, making accurate estimation of higher-order geometric quantities like curvature challenging without additional post-processing. Similarly, the neuronal mitochondria dataset analyzed in this study was previously reconstructed by Mendelsohn *et al.* through extensive manual tracing of each membrane contour slice by slice.^49^ While this approach produced detailed three-dimensional membrane reconstructions, it required substantial subjective human effort, which is difficult to scale. In our framework, manual intervention is currently limited to sparse identification of inside/outside point signals on a small number of slices, substantially reducing annotation effort. In the future, this remaining manual annotation step could be automated by integrating methods such as CNN-based semantic segmentation algorithms.^59,60^ Such integration would further improve the scalability of the frame-work and strengthen its applicability to large-scale experimental datasets.

### E.3 3D membrane curvature heterogeneity in endocytic pits

Our method enabled three-dimensional analysis of curvature patterns in endocytic pits. The reconstructed pits showed azimuthal heterogeneity in curvature around the neck region, where both mean and Gaussian curvature varied around the neck circumference (Fig. 7). This observation is consistent with recent predictions from biophysical simulation studies on membrane budding and neck constriction. Auddya et al. [35] predicted that membrane budding can undergo symmetry-breaking instabilities and transition from axisymmetric to non-axisymmetric morphologies during neck constriction. Their three-dimensional membrane mechanics model suggested that, under biologically relevant membrane tension and curvature-inducing forces, the neck region can spontaneously develop azimuthally heterogeneous curvature as a lower-energy mechanical state. Furthermore, Vasan et al. [36] showed that such non-axisymmetric buckling can lower the energy barrier associated with membrane neck constriction, suggesting that symmetry breaking may play a mechanically favorable role during neck maturation and scission. Our preliminary observations from experimental data are consistent with these theoretical predictions, although further validation using larger datasets is needed. In particular, this measured anisotropy of the mean and Gaussian curvatures in the necks of intermediate- and late-stage endocytic pits may correspond to a lower-energy “shortcut” toward scission of the neck and progression of endocytosis, as predicted by the aforementioned theoretical models. Because many intracellular membrane trafficking processes—including vesicle budding, organelle biogenesis, and membrane fission—also involve highly curved membrane necks, similar symmetry-breaking curvature patterns may represent a more general mechanical principle of membrane remodeling. Furthermore, the reconstructed curvature-resolved membrane geometries provide a foundation for future biophysical modeling based on experimentally measured membrane shapes. Such models could directly evaluate how local curvature distributions, membrane tension, and curvature-generating proteins contribute to neck constriction, scission, or stalled endocytic intermediates.

It should be noted that the current analysis is limited to five reconstructed endocytic pits, and the observed curvature heterogeneity should therefore be regarded as a preliminary, proof-of-concept finding rather than a definitive biological conclusion. As the primary objective of this study is to demonstrate the applicability of the proposed framework for curvature-resolved membrane analysis, establishing statistically robust biological trends is beyond the scope of the present work. Future efforts will extend this analysis to larger populations of endocytic pits, enabling more systematic and statistically powered investigation of three-dimensional curvature patterns during pit morphogenesis.

### E.4 Scope, limitations, and future extensions of physics-informed membrane reconstruction

Although our framework incorporates Helfrich-based physics regularization, it is not restricted to reconstructing smooth membrane geometries or equilibrium configurations that minimize the Helfrich energy. The physics loss acts as a soft constraint rather than a strict requirement, allowing the reconstructed geometry to deviate from the configurations favored by the underlying physical model when supported by observational data. This flexibility arises from the combined optimization of data-fitting and physics-based losses, whose relative contributions can be controlled through adjustable loss weights. Indeed, our framework successfully reconstructed membrane configurations generated using FreeDTS that exhibit thermal fluctuations and protein-induced deformations and do not necessarily correspond to minima of the Helfrich energy (Fig. 2–4). Furthermore, our reconstruction of experimentally observed ER–PM contact sites captured pronounced localized curvature peaks associated with protein-mediated membrane tethering (Fig. 6). These results demonstrate that the proposed physics-informed regularization does not inherently prevent the reconstruction of complex, heterogeneous, or nonequilibrium membrane geometries.

Nevertheless, a fundamental challenge arises when imaging noise and genuine fine-scale membrane features occur at comparable spatial scales and amplitudes. Under such conditions, distinguishing noise from genuine membrane fluctuations becomes inherently difficult, potentially leading to excessive smoothing and the loss of real membrane deformations. This challenge is not specific to PINNs but is common to image-based reconstruction methods that infer geometric structures from noisy or incomplete observations. In our framework, the preservation of fine-scale membrane features depends on their representation in the input point signals and the balance between data fidelity and regularization. Therefore, although the INR provides sufficient representational flexibility to describe complex membrane geometries, the accuracy of the reconstructed features ultimately depends on the quality of the observational data and the extracted point signals.

Another challenge in membrane reconstruction arises from incomplete imaging data, as demonstrated by our missing-wedge analysis (Fig. S7). Incomplete observations make it increasingly difficult to reliably extract both edge/bulk and inside/outside point signals, thereby limiting reconstruction accuracy. Our current framework is not intended to recover missing image information, but rather to reconstruct membrane geometry and curvature from sufficiently well-resolved observational data. Recent studies have explored the use of INRs for missing-wedge recovery,^57^ suggesting a promising direction for integrating image restoration with our physics-informed reconstruction framework.

A separate limitation concerns the representation of complex membrane topologies. Our current phase-field formulation uses only two “colors” {1, −1} to distinguish opposing cellular compartments. Although this binary representation is sufficient for many membrane geometries, biological structures can contain three-compartment and higher-order junctions,^61^ where consistent assignment of inside/outside signals becomes challenging. Extending our framework to such configurations would require modifications to the phase-field representation and associated loss functions, potentially through the use of multiphase implicit functions.^62^

An alluring prospect enabled by this differentiable pipeline is the possibility of using the derived gradients as metrics to estimate the extent of uncertainty in the segmentation process. These uncertainty of the segmented geometries can be propagated into downstream biophysical models of cell signaling such as those enabled by VCell^63^ and SMART.^64^

The use of automatic differentiation and phase-field membrane mechanics stimulates the idea that it could be possible to infer the local material properties of the membrane via optimization. Our current approach is incompatible with this idea owing to a lack of parameter identifiability. A material parameter of interest such as the membrane bending rigidity, *κ_b_*, is absorbed into the _physics_ weighting factor. Additional work, beyond this initial proof of principle will be required to modify the formulation such that the material parameters remain identifiable. Related inference approaches of physical laws and material properties in macroscopic systems have been achieved and are a source of inspiration.^65,66^

## F Conclusion

We present a physics-informed neural network framework for reconstructing membrane geometry from volumetric electron microscopy data using an implicit neural representation. By representing the membrane as a continuous phase field and incorporating physics-based regularization, our approach enables accurate reconstruction of membrane morphology, particularly in higher-order differential geometric quantities such as mean and Gaussian curvatures. Overall, this work bridges image-based membrane reconstruction with quantitative biophysical analysis by enabling curvature-resolved characterization of membrane geometry in three dimensions. This capability provides a foundation for probing the physical principles underlying membrane shape and for connecting geometric features to underlying biological mechanisms.

## G Methods

This section presents the mathematical formulation of the proposed membrane reconstruction framework. Descriptions of all model parameters and their values used in this study are summarized in Table 2.

**Table 2:** Model parameters. The values listed here were used throughout this study unless otherwise specified. *Σ* denotes the half-length of the normalized computational domain of the implicit neural field, defined as [−*σ, σ*]^3^.

| Symbol | Value | Unit | Description |
|---|---|---|---|
| *n*<sub>layer</sub> | 2 | - | Number of hidden layers |
| *n*<sub>hidden</sub> | 128 | - | Number of neurons per layer |
| *ϵ* | 0.05 | *σ* | Thickness of membrane |
| *τ*<sub>mem</sub> | 0.75 | - | Threshold for membrane (edge) region |
| *τ*<sub>non</sub> | 0.75 | - | Threshold for non-membrane (bulk) region |
| *τ*<sub>smooth</sub> | 0.02 | - | Threshold for smoothing loss collocation points |
| *λ*<sub>data</sub> | 100,000 | - | Weight for data loss |
| *λ*<sub>B.C.</sub> | 100,000 | - | Weight for boundary loss |
| *λ*<sub>physics</sub> | 1 | - | Weight for physics loss |
| *λ*<sub>smooth</sub> | 10 | - | Weight for smoothing loss |
| *N*<sub>physics</sub> | 80,000 | - | Number of collocation points for physics loss |
| *N*<sub>smooth</sub> | 5,000 | - | Number of collocation points for smoothing loss |
| *η* | 0.001 | - | Learning rate of optimization |
| *N*<sub>iter</sub> | 10,000 | - | Number of optimization iterations per each stage |
| *N*<sub>init</sub> | 5,000 | - | Number of iterations to set initial conditions |

### G.1 Implicit neural representation of membrane

We represent the membrane geometry using phase field *φ* : Ω R, which takes *φ* = 1 in the bulk regions and whose zero level set defines the membrane surface, = **x** Ω *φ*(**x**) = 0.^44–46^ The continuous phase field is parameterized by a neural network with parameters *θ*, yielding an implicit neural representation (INR) *φ_θ_* : Ω R.^20,24,25^ The network takes spatial coordinates **x** Ω as input and outputs a scalar value *φ_θ_*(**x**), representing the phase-field value at that location.

The network is implemented as a fully connected MLP with *N*_layers_ hidden layers and *N*_hidden_ neurons per layer. Each layer consists of an affine transformation followed by a nonlinear activation; the hyperbolic tangent function, tanh(·), is used as the activation throughout.

As a coordinate-based representation, *φ_θ_* defines a continuous and differentiable function over the spatial domain, enabling evaluation at arbitrary spatial locations independent of the underlying voxel grid from the three-dimensional image data. This property allows spatial derivatives of *φ_θ_* to be computed via automatic differentiation, which is essential for formulating physics-based loss functions and for computing surface normals and curvatures. Notably, implicit neural representations can capture fine geometric details and complex surface structures without being constrained by a fixed spatial resolution.

### G.2 Loss functions

#### G.2.a Enforcing consistency with the image (data and boundary losses)

To satisfy the first requirement—that the reconstructed membrane geometry be consistent with the volumetric image data—we define data loss, _data_, and boundary loss, _B.C._. These loss functions constrain the reconstructed membrane phase field, *φ_θ_*, to match the membrane surface inferred from the image intensity values, *I*(**x***_i_*).

To compute the data loss, we first sample point signals for membrane and non-membrane regions. For this proof of principle, we identify these regions using simple intensity thresholding, although the labels could alternatively be obtained from semantic segmentation algorithms.^16–19,58^ Let *τ*_mem_ and *τ*_non_ denote the membrane and non-membrane intensity thresholds, respectively.

Membrane and non-membrane signal points are randomly sampled from voxel centers satisfying _mem_ **x***_j_* Ω *I*(**x***_j_*) *τ*_mem_ and _non_ **x***_j_* Ω *I*(**x***_j_*) *τ*_non_. The data set is then defined as _data_ = _mem non_, where each point **x***_j_* _data_ is assigned a binary label *y*(**x***_j_*) 0, 1, with *y* = 1 for membrane points and *y* = 0 for non-membrane points. For convenience, we refer to these binary labels as edge/bulk signals throughout the manuscript.

Next, we define the membrane energy density

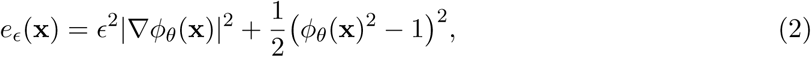

where *E >* 0 controls the characteristic width of the diffuse interface. The energy density acts as a diffuse surface indicator: it is localized near the membrane interface and approac<u>h</u>es zero in the bulk. When the phase field approximates the equilibrium profile 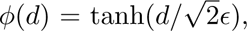, where *d* is the signed distance from the membrane surface, *e_E_* attains a unit maximum at the interface (*φ* = 0).

Using this energy density, we define the data loss as

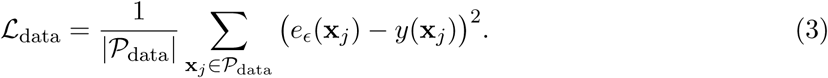

Minimizing this loss aligns the zero level set *φ_θ_* = 0 with membrane signals while promoting a locally equilibrated tanh-like phase field profile across the membrane interface.

Although the data loss stabilizes the phase field in the bulk by favoring *φ* 1, it is symmetric with respect to the sign of *φ* and therefore cannot determine which side of the membrane corresponds to *φ* = +1 or *φ* = 1. To resolve this ambiguity, we introduce a boundary loss that enforces consistent phase labeling throughout the bulk.

To compute the boundary loss, we sample spatial locations in bulk regions and assign target signs. Specifically, we randomly sample a set of coordinates _bulk_ Ω from regions expected to lie far from the membrane surface. Each sampled point **x***_j_* _bulk_ is assigned a sign label *s*(**x***_j_*) +1, 1 indicating the side of the membrane on which it lies. The sign labels may be assigned manually or inferred from image cues, such as intensity differences between the membrane interior and exterior, or obtained using semantic segmentation algorithms.^16–19,58^ Throughout the manuscript, we refer to these labels as inside/outside signals.

Using these bulk sign points, the boundary loss is defined as

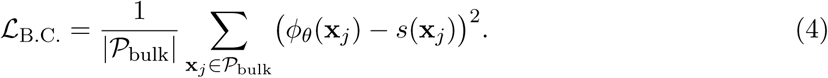

Minimizing _B.C._ breaks the sign symmetry of the phase field, suppressing spurious sign inversions away from the membrane surface.

#### G.2.b Recovering physically consistent curvature (physics and smoothing losses)

Volumetric image data provide information about membrane shape but only weakly constrain its curvature distribution. This is because curvature depends on higher-order derivatives of the membrane surface (Fig. 1A), which cannot be accurately recovered from a discrete voxel representation, particularly in the presence of imaging noise. Consequently, the data and boundary losses alone are insufficient to recover a physically consistent curvature distribution. To satisfy the second requirement—that the reconstructed membrane geometry recover a physically consistent curvature distribution—we define the physics loss, _physics_, and smoothing loss, _smooth_.

The physics loss is based on the Helfrich-Canham-Evans bending energy, the most widely used continuum model of membrane mechanics.^29–31^ This model describes the elastic cost of bending a lipid bilayer by penalizing membrane curvature.^33,67^ For vanishing spontaneous curvature, the bending energy is given by *E_b_* = 2*κ H*^2^*dA*, where *H* is the mean curvature, *κ* is the bending rigidity, and the integral is taken over the membrane surface. In the phase-field formulation, this energy is expressed as^20,44–46^

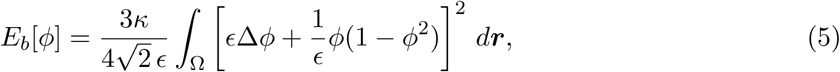

where *E* again controls the diffuse interface thickness. Biological membranes tend to adopt configurations that minimize this energy functional.

To incorporate the Helfrich bending energy into the optimization, we approximate the volumetric integral by Monte Carlo sampling over randomly selected collocation points. We first sample *N*_physics_ collocation points, denoted by P_physics_ ⊂ Ω, throughout the spatial domain. At each collocation point **x***_j_* ∈ P_physics_, we evaluate

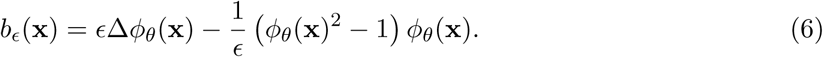

The physics loss is then defined as

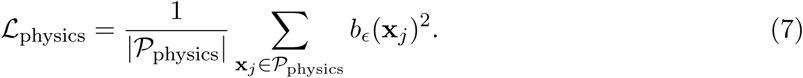

Minimizing _physics_ drives the reconstructed membrane toward configurations with lower Helfrich bending energy, thereby regularizing the curvature distribution toward physically realistic solutions. To further suppress spurious fluctuations in the curvature distribution and obtain a smooth, physically realistic solution, we introduce a smoothing loss that directly penalizes variations in curvature along the membrane surface. We first sample *N*_smooth_ collocation points for smoothing loss evaluation from the zero level set of the phase field, *P*_smooth_ **x***_j_* Ω *φ_θ_*(**x***_j_*) *< τ*_smooth_, where *τ*_smooth_ *>* 0 is a threshold that defines a narrow band around the zero level set of the phase field. As described in the next section, the smoothing loss is applied only during the third stage of the optimization. Accordingly, this sampling is performed after completion of the second stage.

The smoothing loss is defined as

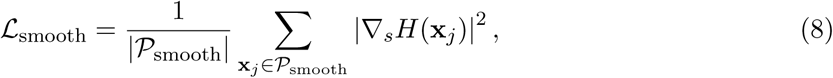

where *_s_* represents the surface gradient operator and *H* denotes the mean curvature, computed by Eq. (11) and Eq. (12), respectively.

### G.3 Model training

#### G.3.a Optimization and implementation

Optimization is performed using the Adam optimizer with learning rate *η*. Training is carried out in three stages, each consisting of *N*_iter_ optimization steps. In the first stage, the weights for the physics and smoothing losses are set to zero (*λ*_physics_ = 0, *λ*_smooth_ = 0), and the network was optimized using only the data and boundary loss terms. In the second stage, the smoothing loss remains inactive (*λ*_smooth_ = 0), and optimization is performed using the data, boundary, and physics loss terms. In the third stage, all loss terms are activated, including the data, boundary, physics, and smoothing losses.

The optimization is implemented in JAX,^68^ leveraging automatic differentiation to compute first- and second-order spatial derivatives of the phase field. All computations are performed using GPU acceleration.

#### G.3.b Network initialization

At the beginning of the first stage, the network is initialized to represent a spatially uniform phase field. This is achieved by performing an initialization phase consisting of *N*_init_ steps, during which the following loss function is minimized.

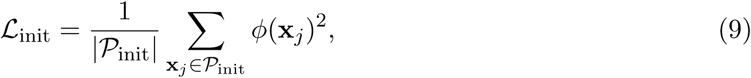

where *P*_init_ denotes a set of points randomly sampled from the simulation domain. This initialization biases the network toward *φ* = 0 everywhere prior to the main optimization, providing a neutral starting point without introducing spurious geometric structure.

#### G.3.c Hyperparameter selection

The loss weights were selected empirically to balance the contributions of the individual loss terms. Because the loss terms have different numerical scales, we monitored their magnitudes during optimization and adjusted the weights such that the weighted loss contributions, *λ_i i_*, were of comparable orders of magnitude. This served as a practical guideline rather than a strict requirement, with the final parameter values determined based on optimization stability and reconstruction accuracy.

### G.4 Post-processing

#### G.4.a Surface extraction

After optimization, a continuous phase field *φ_θ_*(**x**) is obtained over the three-dimensional simulation domain. The membrane surface is defined as the zero level set of the phase field, *φ_θ_* = 0. To extract an explicit surface representation, we apply the marching cubes algorithm^69^ to the discretized phase field, yielding a triangular mesh approximation of the *φ_θ_*= 0 isosurface.

#### G.4.b Computation of geometric quantities

Various geometric quantities associated with the membrane are computed from the phase field, *φ_θ_*(**x**), using JAX’s automatic differentiation framework.^68^ These quantities are evaluated on the zero level set of *φ_θ_*(**x**).

##### G.4.b.1 Surface normal

The unit normal vector to the membrane interface is obtained by normalizing the gradient of the phase field,

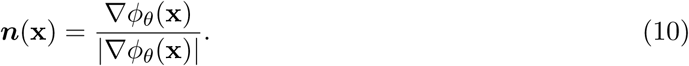

##### G.4.b.2 Surface gradient operator

The surface gradient operator is defined as the projection of the Euclidean gradient onto the local tangent plane,

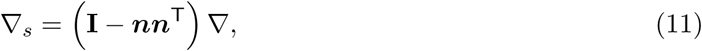

where **I** is the identity matrix.

##### G.4.b.3 Mean curvature

The mean curvature is computed as one-half of the divergence of the unit normal vector. Expressing the normal in terms of the phase field yields

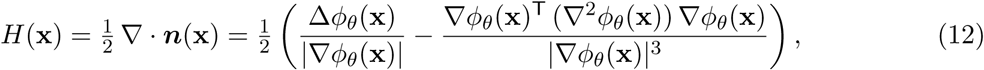

where ∇^2^*φ_θ_* denotes the Hessian matrix of *φ_θ_*and Δ*φ_θ_* = tr(∇^2^*φ_θ_*) denotes the Laplacian.

##### G.4.b.4 Gaussian curvature

The Gaussian curvature is computed from the gradient and Hessian of the phase field according to

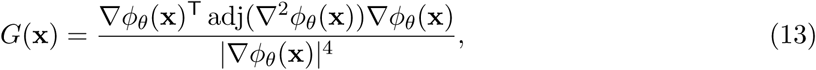

where ∇^2^*φ_θ_* denotes the Hessian matrix of *φ_θ_*and adj(·) denotes the adjugate matrix.

### G.5 Analysis

#### G.5.a Surface Dice score

To quantify the agreement between the reconstructed membrane surfaces and the ground-truth geometries of synthetic datasets or reference segmentations of experimental datasets, we calculated the surface Dice score.^70^ For each surface, we defined a thin representation of the membrane surface and a surrounding region within a specified distance tolerance. We then evaluated surface agreement bidirectionally, from the reconstructed surface to the reference surface and vice versa, corresponding to precision and recall, respectively. Precision was defined as the fraction of reconstructed surface points lying within the specified distance tolerance of the reference surface, while recall was defined as the fraction of reference surface points lying within the same tolerance of the reconstructed surface. The surface Dice score was computed as the harmonic mean of precision and recall,

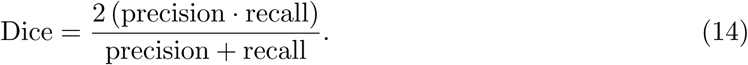

A Dice score of 1 indicates perfect agreement between the reconstructed and reference surfaces, while a score of 0 indicates no surface agreement within the specified distance tolerance.

#### G.5.b Geometric correlation analysis

To quantify the accuracy of reconstructed differential geometric quantities, we evaluated the agreement between the reconstructed membrane and the ground truth in terms of normal angle, mean curvature, and Gaussian curvature. We first extracted the reconstructed membrane surface as the zero level set of the phase field using the marching cubes algorithm and sampled points on this surface. For each sampled point, we identified the closest point on the ground truth surface using a nearest-neighbor search, thereby establishing a pointwise correspondence between the reconstructed and ground truth surfaces. At each matched pair of points, we compared the corresponding scalar geometric quantities, namely normal angle, mean curvature, and Gaussian curvature. The normal angle was defined as the angle between the local surface normal and a fixed vertical reference direction. To obtain a global measure of agreement, we computed Pearson’s correlation coefficient between the reconstructed and ground truth values across all sampled points for each quantity. A higher correlation indicates stronger agreement in the spatial distribution of the corresponding geometric feature.

#### G.5.c Surface distance metrics

To quantify the geometric differences between the reconstructed and reference membrane surfaces, we calculated the average surface distance (ASD) and Hausdorff distance (HD). For each point on the reconstructed surface, we computed the Euclidean distance to the nearest point on the reference surface, and repeated this calculation in the opposite direction. The ASD was calculated by averaging the nearest-surface distances in both directions, whereas HD was defined as the maximum of these distances. We also calculated the 95th percentile Hausdorff distance (HD95) to reduce sensitivity to extreme outliers.

### G.6 Experimental data processing

#### G.6.a Boundary treatment and exclusion of unreliable regions

For experimental datasets, incomplete observations and image boundaries can introduce ambiguities in the extraction of edge/bulk and inside/outside point signals. To mitigate these effects, we manually excluded image planes where reliable point signals could not be extracted prior to PINN reconstruction. Following reconstruction, we visually compared the reconstructed membrane geometry with the original volumetric images and manually cropped regions showing discrepancies or unreliable reconstruction, excluding them from subsequent curvature analysis.

#### G.6.b Extraction of the neck ring of endocytic pits

To quantify neck geometry of endocytic pits, we extracted a representative neck ring from the reconstructed implicit membrane surface. First, a neck band was identified as the largest connected component of vertices exhibiting strongly negative Gaussian curvature. A local neck plane was then estimated by applying principal component analysis to the neckband vertices, with the first two principal directions defining the plane and the third defining its normal. The phase field was sampled on this plane, and the membrane contour was obtained as the zero level set (*φ_θ_*= 0). Among the extracted contours, the one corresponding to the neck band was selected and mapped back into three-dimensional space to obtain the neck ring for subsequent geometric analysis.

### G.7 Datasets

#### G.7.a Synthetic volume image data

##### G.7.a.1 Mesh generation

We first generated membrane geometries as triangulated meshes using the membrane simulator, Mem3DG^39^ and FreeDTS^40^ (Fig. S1). Both simulators model membrane using Helfrich energy, making them well suited for the analysis of present study, as the proposed framework is designed to reconstruct membrane surfaces and infer higher-order geometric quantities under the same physical principles.

To generate the biconcave disk geometry, we used Mem3DG. Starting from an oblate spheroidal membrane with zero spontaneous curvature, we simulated membrane deformation under prescribed preferred surface area and enclosed volume, resulting in a biconcave morphology at equilibrium. To generate the Ω-shaped bud geometry, we also used Mem3DG. An initially planar membrane with open boundary conditions was subjected to a localized spontaneous curvature, which induced membrane invagination and formed a budded morphology with a narrow neck. To generate the asymmetric protrusion geometry, we used FreeDTS. Starting from an initially planar membrane with periodic boundary conditions, we introduced curvature-inducing proteins covering 20% of the membrane surface. Thermal fluctuations and attractive protein–protein interactions promoted protein clustering and local membrane deformation, resulting in asymmetric membrane protrusions. Surface normals, mean curvature, and Gaussian curvature were computed from the resulting membrane meshes using Mem3DG and served as ground truth for quantitative evaluation of the reconstruction framework.

##### G.7.a.2 Volumetric image generation

We generated synthetic volumetric imaging data from the triangulated surface meshes to serve as the data constraints for our learning framework. For each geometry, the final mesh was embedded in a uniform Cartesian grid of size 128 128 128 and voxelized to generate a binary membrane mask. A Euclidean distance transform was then applied to generate a membrane-like intensity profile, such that voxels near the membrane exhibited high intensity, while voxels farther away decayed exponentially in intensity. To mimic experimental image degradation, random noise, both additive and subtractive, was introduced into the voxelized membrane masks prior to distance transformation. Specifically, a prescribed fraction of membrane voxels was randomly flipped to background voxels, while a prescribed fraction of background voxels was randomly flipped to membrane voxels. The fraction of perturbed voxels was systematically varied from 0.0 to 0.8 in the ablation study and fixed at 0.6 in the main analyses. The resulting image volumes were smoothed using a Gaussian filter and normalized such that the maximum intensity was one.

Using the generated volumetric data, edge and bulk signals were extracted by the intensity-based thresholding procedure of the framework. Inside and outside signals were randomly sampled based on the ground truth geometry.

#### G.7.b Experimental tomography data

##### G.7.b.1 Golgi apparatus

Publicly available cryo-electron tomograms of the Golgi apparatus were downloaded from the Chan Zuckerberg Imaging Institute CryoET Data Portal.^50^ We used the dataset reported by Khavnekar et al.^48^ (Dataset ID: DS-10301, Deposition ID: CZCDP-10300, Run ID: RN-14085). The dataset consists of in situ cryo-electron tomography images of *Chlamydomonas reinhardtii*, cryogenically preserved and prepared by cryo-plasma focused ion beam milling, and reconstructed at a voxel size of 0.784 nm.^48^

The tomograms were preprocessed by cropping sub-volumes containing the Golgi apparatus, enhancing the contrast between membrane and non-membrane regions using contrast-limited adaptive histogram equalization (CLAHE),^71^ and normalizing the voxel intensities to the range [0, 1]. From the preprocessed tomograms, edge and bulk signals were extracted using a simple intensity-thresholding method, while inside and outside signals were assigned by manual inspection of the membrane geometry.

##### G.7.b.2 Mitochondria

Three-dimensional electron tomograms of mitochondria were used from the dataset by Mendelsohn et al..^49^ The dataset consists of serial-section electron tomography images of neuronal mitochondria in mouse cerebellum neuropil, chemically preserved by high-pressure freezing followed by freeze substitution and resin embedding, and acquired at an isotropic voxel size of 1.64 nm.^49^

We first cropped sub-volumes containing mitochondria and roughly isolated the mitochondrial regions from the surrounding structures using Segment Anything Model 3.^59^ The resulting volumes were then preprocessed by enhancing the image contrast using CLAHE^71^ and normalizing the voxel intensities to the range [0, 1]. From the preprocessed tomograms, edge and bulk signals were extracted using a simple intensity-thresholding method, while inside and outside signals were assigned by manual inspection of the membrane geometry.

##### G.7.b.3 ER–plasma membrane contact sites

Cryo-electron tomography data of endoplasmic reticulum (ER)–plasma membrane (PM) contact sites were downloaded from Collado et al.^51^ (EMDB ID: EMD-10378). The dataset consists of an in situ cryo-electron tomogram of a heat-shocked wild-type *Saccharomyces cerevisiae* cell, cryogenically preserved and prepared by cryo-focused ion beam milling, and reconstructed at a voxel size of 1.368 nm. The tomogram captures cortical ER membranes adjacent to the PM, including regions of high membrane curvature associated with ER–PM tethering.

The tomogram was preprocessed by cropping a sub-volume containing the ER–PM contact site. We then applied MemBrain-seg^16^ to segment the membranes and used the resulting segmentation to sample edge/bulk and inside/outside point signals.

##### G.7.b.4 Clathrin-mediated endocytic pits

Three-dimensional electron microscopy data of clathrin-mediated endocytic pits were downloaded from the publicly available OpenOrganelle portal^72^ pro-vided by HHMI Janelia Research Campus. We used dataset reported by Xu et al.^14,52^ (Dataset ID: jrc_cos7-1b). The dataset consists of FIB-SEM images of an interphase COS-7 cell, chemically preserved by high-pressure freezing followed by freeze substitution, heavy-metal staining, and resin embedding, and reconstructed with a voxel size of 2 nm.^14,52^

We first identified membrane invaginations consistent with clathrin-mediated endocytic pits by manual inspection of the volumetric data and cropped 5 sub-volumes containing individual endocytic structures. The cropped volumes were then preprocessed by enhancing the image contrast using CLAHE^71^ and normalizing the voxel intensities to the range [0, 1]. From the preprocessed tomograms, edge and bulk signals were extracted using a simple intensity-thresholding method, while inside and outside signals were assigned by manual inspection of the membrane geometry.

## H Source Code

The source code used for all work in this study is hosted on https://github.com/ctleelab/pinn-exploration-model and archived on Zenodo.^73^

## I Author Contributions

Conceptualization: AM, MA, CTL; Data curation: AM; Formal analysis: AM, SK; Funding Acquisition: MA, CTL; Investigation: AM, SK; Methodology: AM, CTL; Project administration: MA, CTL; Resources: MA, CTL; Software: AM, SK, CTL; Supervision: MA, CTL; Validation: AM; Visualization: AM, SK; Writing-original draft: AM, MA, CTL; Writing-review and editing: AM, SK, MA, CTL.

## J Acknowledgments

This work was supported by The Gordon and Betty Moore Foundation’s Postdoctoral Fellowship to AM, NIH R00GM132551 to MA, and NIH R35GM166376 to CTL. This work utilized computational resources provided by the Hyak supercomputer system at the University of Washington. We are also grateful to the UCSD Physics Computing Facility for computational resources.

## Supplemental Material

**Figure S1:**
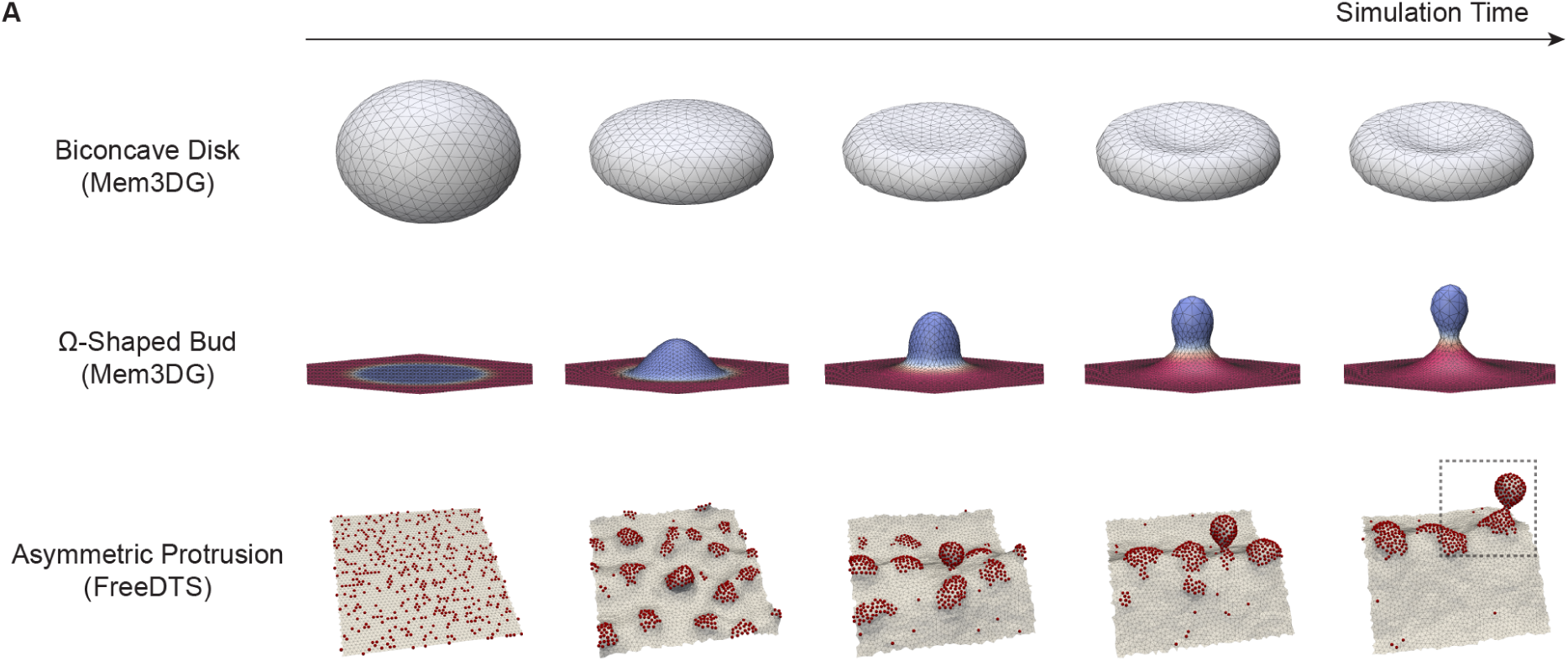
Snapshots of membrane simulations used to generate synthetic datasets. Ground-truth membrane meshes were generated using Mem3DG and FreeDTS. The biconcave disk geometry was obtained by relaxing an initially oblate spheroidal membrane under prescribed preferred surface area and enclosed volume constraints. The Ω-shaped bud geometry was generated by imposing a spatially heterogeneous spontaneous curvature on an initially planar membrane. Red and blue surfaces indicate spontaneous curvatures of *C*0 = 0.0 and 8.0 (unit length)*^−^*^1^, respectively. The asymmetric protrusion geometry was generated by introducing curvature-inducing proteins (red dots) onto a planar membrane, where thermal fluctuations and attractive protein–protein interactions promoted protein clustering and membrane deformation. Dashed boxes indicate the subvolumes selected for PINN reconstruction.

**Figure S2:**
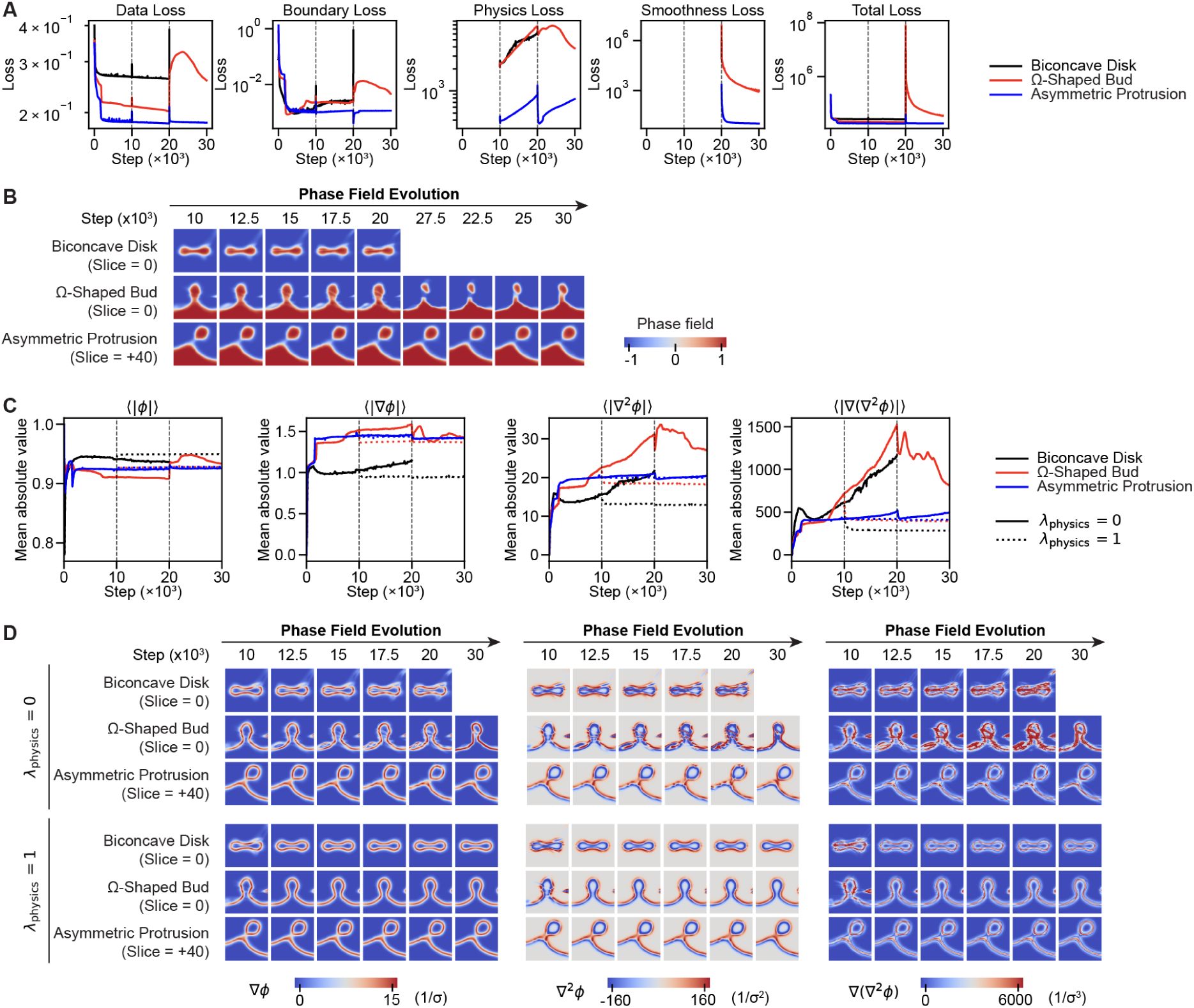
Role of physics regularization. To investigate the role of physics regularization, we applied PINN to synthetic data without the physics loss term (i.e., *λ*_physics_ = 0). The evolution of the loss functions and phase fields during optimization is shown in panels (A) and (B), respectively. The optimization failed for the biconcave disk and Ω-shaped bud, whereas it succeeded for the protein-induced peak. To further understand the effect of physics regularization, we examined the derivative fields, namely ∇*φ*, ∇^2^*φ*, and ∇(∇^2^*φ*), and compared the results obtained with and without physics regularization (*λ*_physics_ = 1 and 0, respectively). Panel (C) shows the spatial ensemble averages of these quantities over the course of optimization, while panel (D) illustrates their spatial distributions. Data are not shown for cases in which simulations failed due to numerical instability.

**Figure S3:**
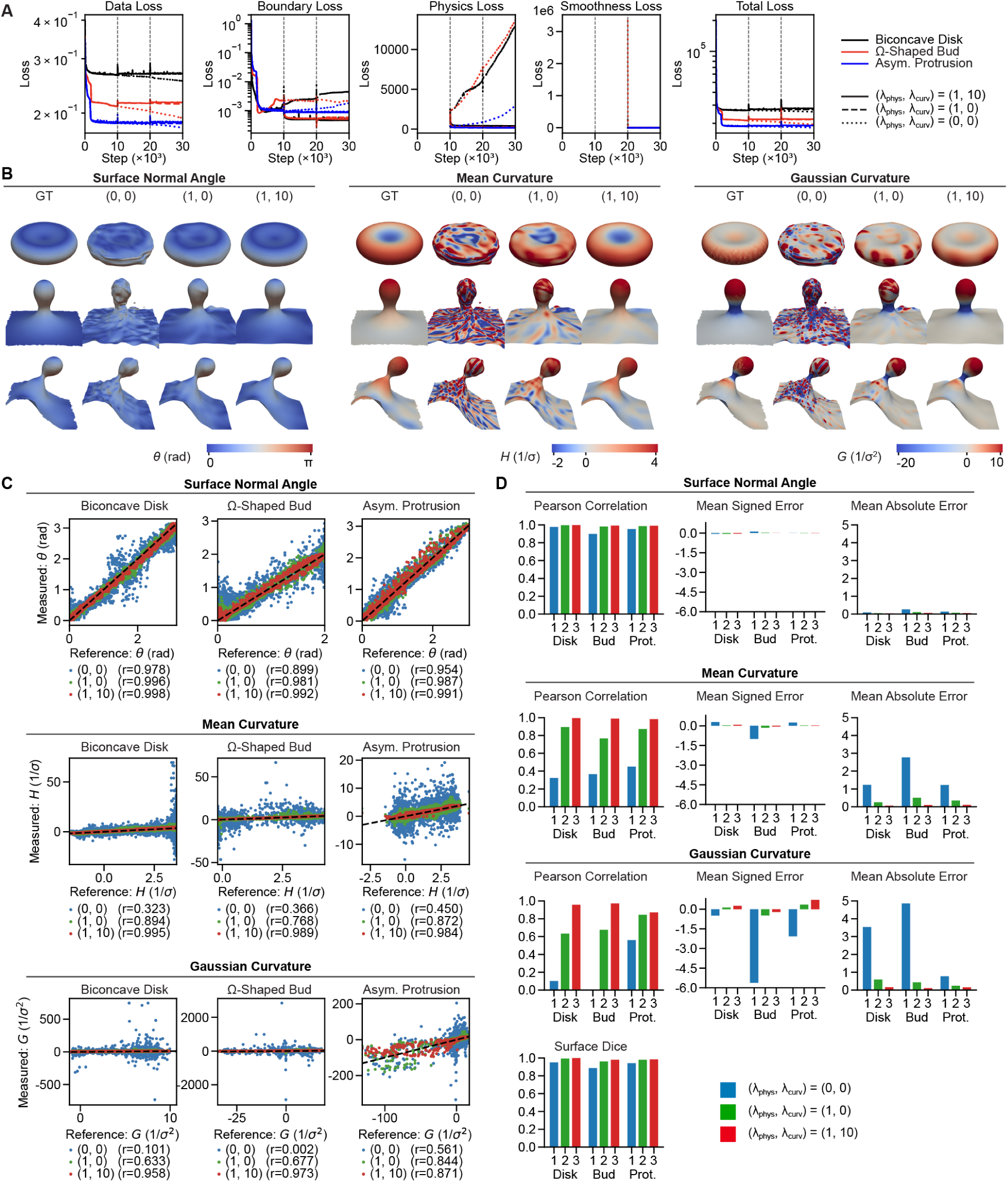
Longer optimization does not compensate for the physics and smoothing losses. To eliminate the effect of optimization length when evaluating the contributions of the physics and smoothing losses in Fig. 3, we extended all optimization runs to a total of 30 × 10^3^ steps. We considered three cases with different loss configurations. In the first case, optimization was continued after Stage 1 (10 × 10^3^ steps) without introducing additional loss terms, maintaining (*λ*_physics_*, λ*_smooth_) = (0, 0) until 30 × 10^3^ steps. In the second case, the physics loss was introduced after Stage 1, with (*λ*_physics_*, λ*_smooth_) = (1, 0), and optimization was continued until 30 × 10^3^. The third case followed the full three-stage optimization procedure, in which the physics loss was introduced at Stage 2 and the smoothing loss was subsequently introduced at Stage 3, with (*λ*_physics_*, λ*_smooth_) = (1, 10). (A) Progression of the loss functions during optimization. Solid, dashed, and dotted lines represent the three-, two-, and one-stage optimization procedures, respectively. (B) Surface properties reconstructed after 30 × 10^3^ optimization steps. The parameter pairs (0, 0), (1, 0), and (1, 10) denote the one-, two-, and three-stage optimization procedures, respectively. (C) Comparison of the reconstructed surface properties with their corresponding ground-truth values. (D) Quantitative evaluation of reconstruction accuracy using correlation, signed error, and absolute error between the reconstructed and ground- truth surface properties.

**Figure S4:**
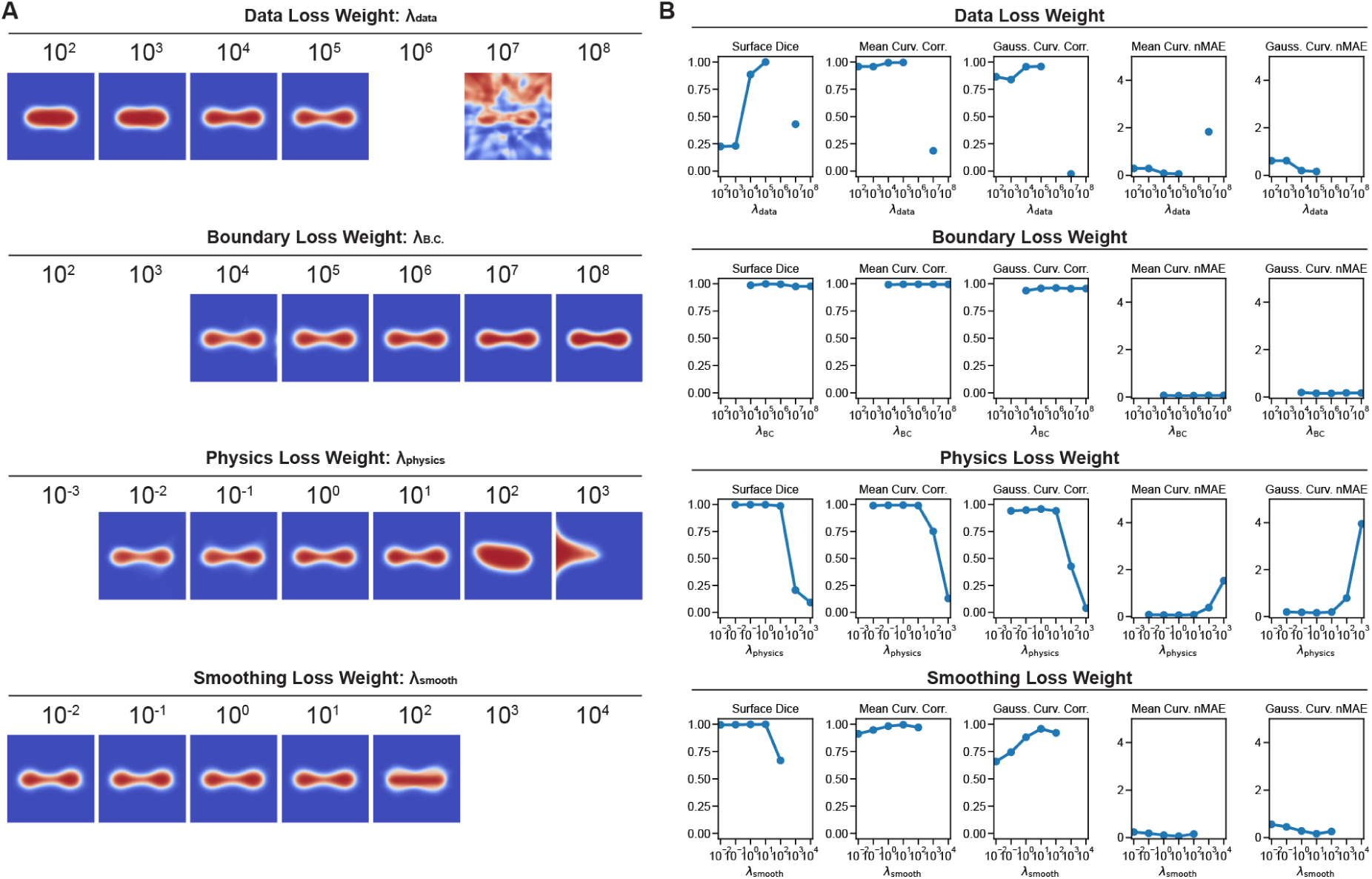
Effect of loss weights. We reconstructed the synthetic biconcave disk geometry by varying each loss weight individually while keeping the remaining weights fixed at their reference values, (*λ*_data_*, λ*_B.C_*., λ*_physics_*, λ*_smooth_) = (10^5^, 10^5^, 10^0^, 10^1^). (A) Mid-plane cross-sections of the reconstructed phase field at the final training step. (B) Quantitative evaluation of surface and curvature reconstruction accuracy using the surface Dice coefficient, Pearson correlation coefficients between reconstructed and ground-truth mean and Gaussian curvatures, and normalized mean absolute errors (nMAEs) of the corresponding curvature estimates. Data are not shown for simulations that failed due to numerical instability.

**Figure S5:**
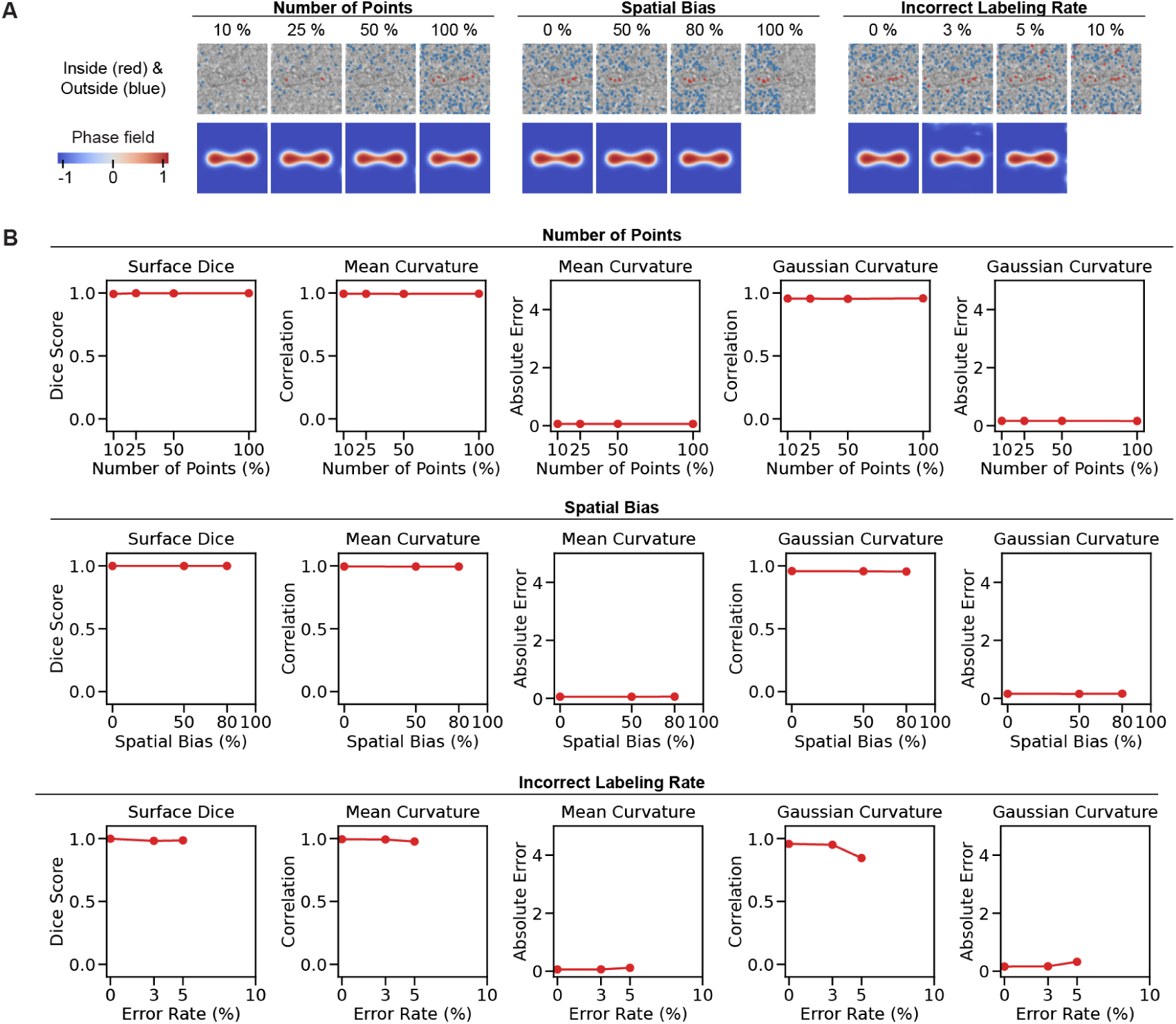
Effect of the choice of inside/outside point signals. We performed membrane reconstructions of a biconcave disk using different configurations of inside/outside point signals. Specifically, we varied the total number of points (with 20,000 points as the reference, corresponding to 100%), their spatial distribution (by introducing a specified degree of sampling bias toward one side along a given axis), and labeling accuracy (by randomly assigning incorrect inside/outside labels at specified error rates). All other point signals were kept constant, including the reference edge/bulk point signals at a noise level of 0.7 and the same set of physics collocation points. (A) Distributions of inside/outside point signals and the corresponding PINN reconstruction results (phase fields) visualized in the midplane. (B) Quantitative evaluation of reconstruction accuracy using surface Dice, Pearson correlation coefficients, and mean absolute errors for mean and Gaussian curvatures. Data are not shown for simulations that failed due to numerical instability.

**Figure S6:**
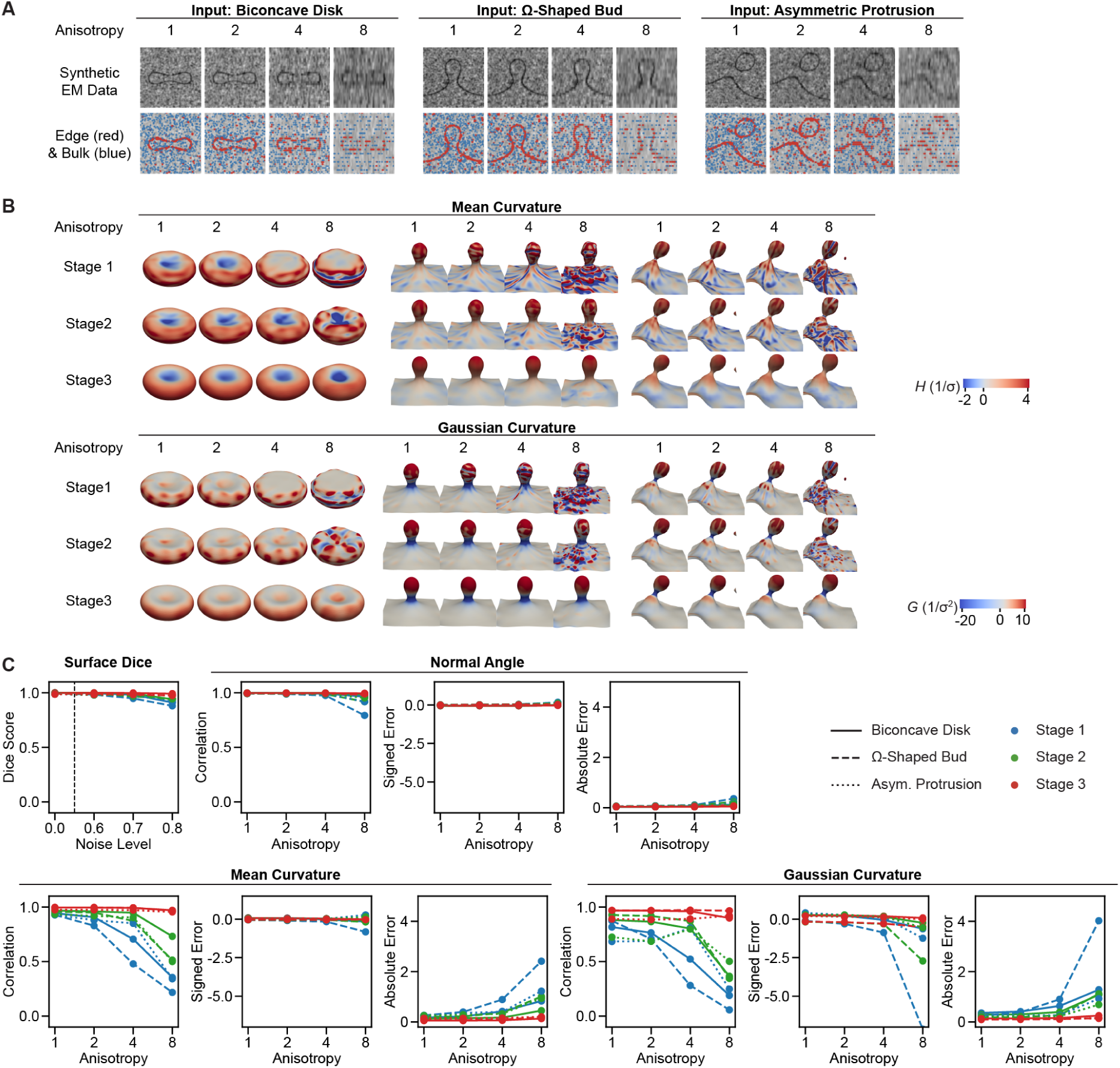
Accuracy of reconstructed geometries under increasing anisotropic voxel resolution. (A) Synthetic EM images and extracted edge/bulk signals with increasing anisotropy. (B) Three-dimensional reconstruction of membrane shape and curvature distributions at each optimization stage across different levels of anisotropy. (C) Quantitative agreement with the ground truth, assessed using the surface Dice score for geometric agreement, as well as Pearson correlation, mean signed error, and mean absolute error for the surface normal angle, mean curvature, and Gaussian curvature.

**Figure S7:**
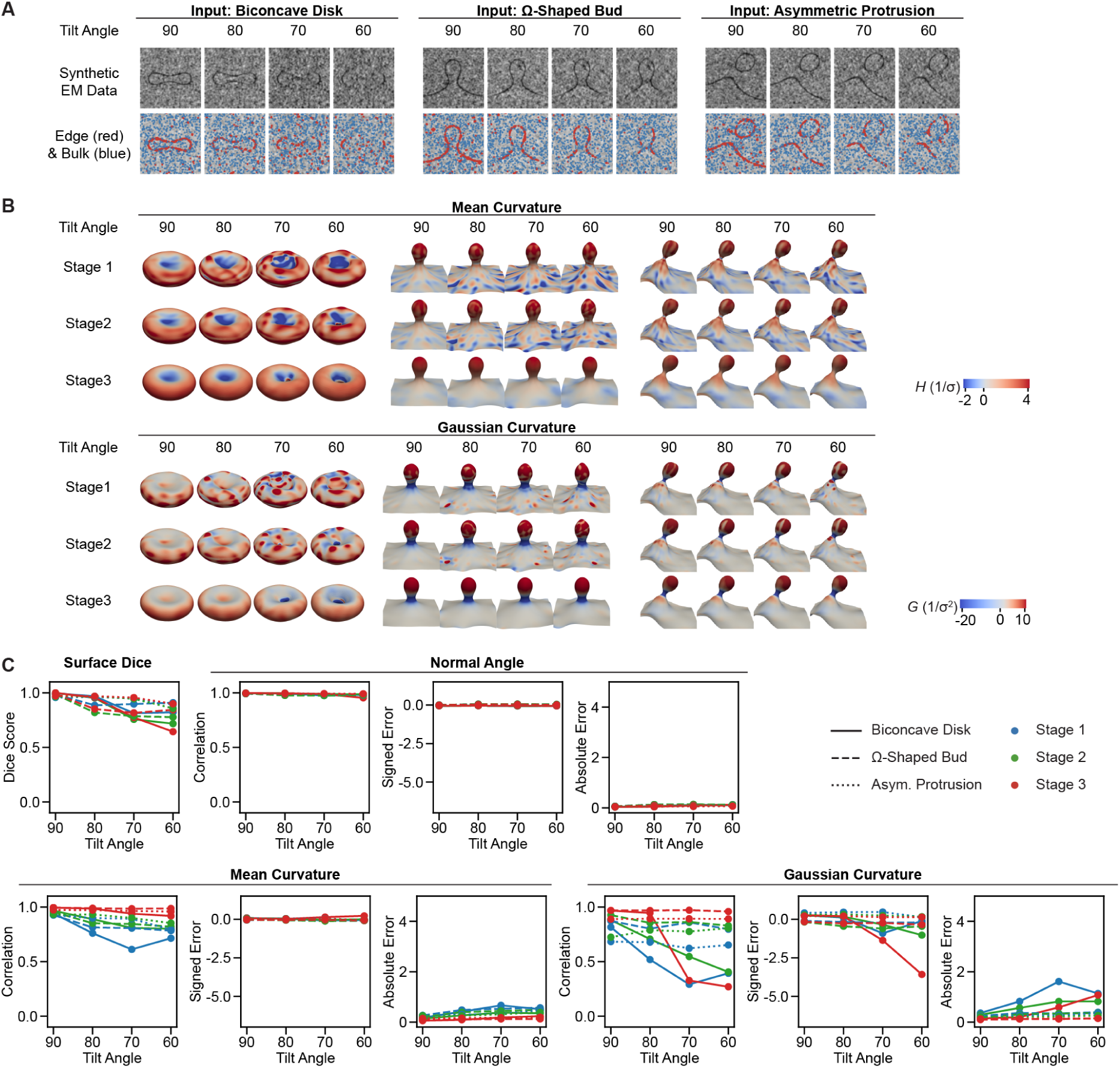
Accuracy of reconstructed geometries with increasing degree of missing wedge. (A) Synthetic EM images and extracted edge/bulk signals with decreasing tilt angles. (B) Three-dimensional reconstruction of membrane shape and curvature distributions at each optimization stage across different tilt angles. (C) Quantitative agreement with the ground truth, assessed using the surface Dice score for geometric agreement, as well as Pearson correlation, mean signed error, and mean absolute error for the surface normal angle, mean curvature, and Gaussian curvature.

**Figure S8:**
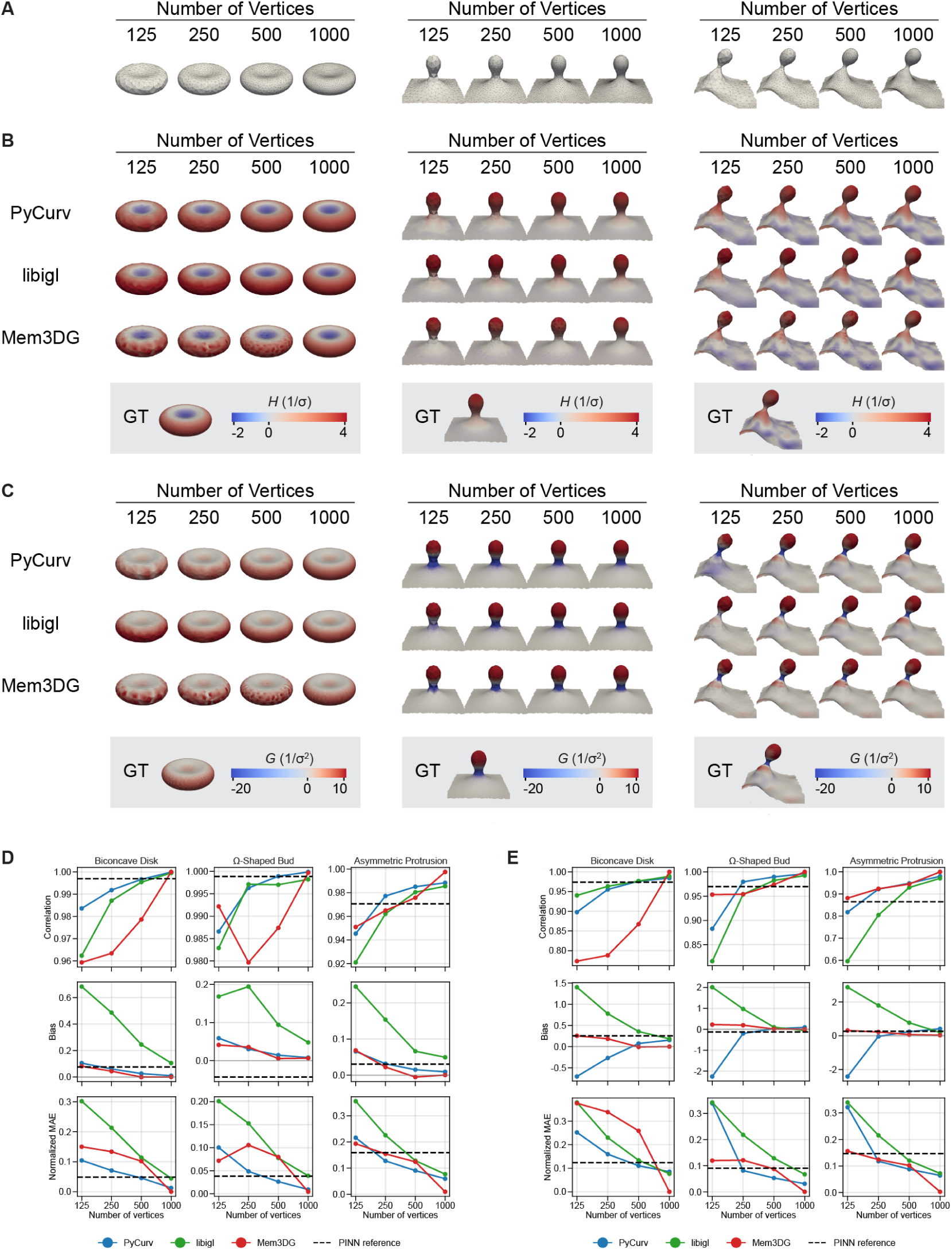
Curvature estimation using mesh-based membrane representations. (A) Synthetic membrane meshes with varying numbers of vertices. (B–C) Mean curvature (B) and Gaussian curvature (C) estimated using three mesh-based methods: PyCurv, libigl, and Mem3DG. (D–E) Accuracy of mean curvature (D) and Gaussian curvature (E) estimates evaluated against ground-truth values using Pearson correlation, mean signed error, and mean absolute error. Solid lines represent the results of mesh-based methods, while dashed lines indicate the corresponding PINN-based estimates for comparison.

**Figure S9:**
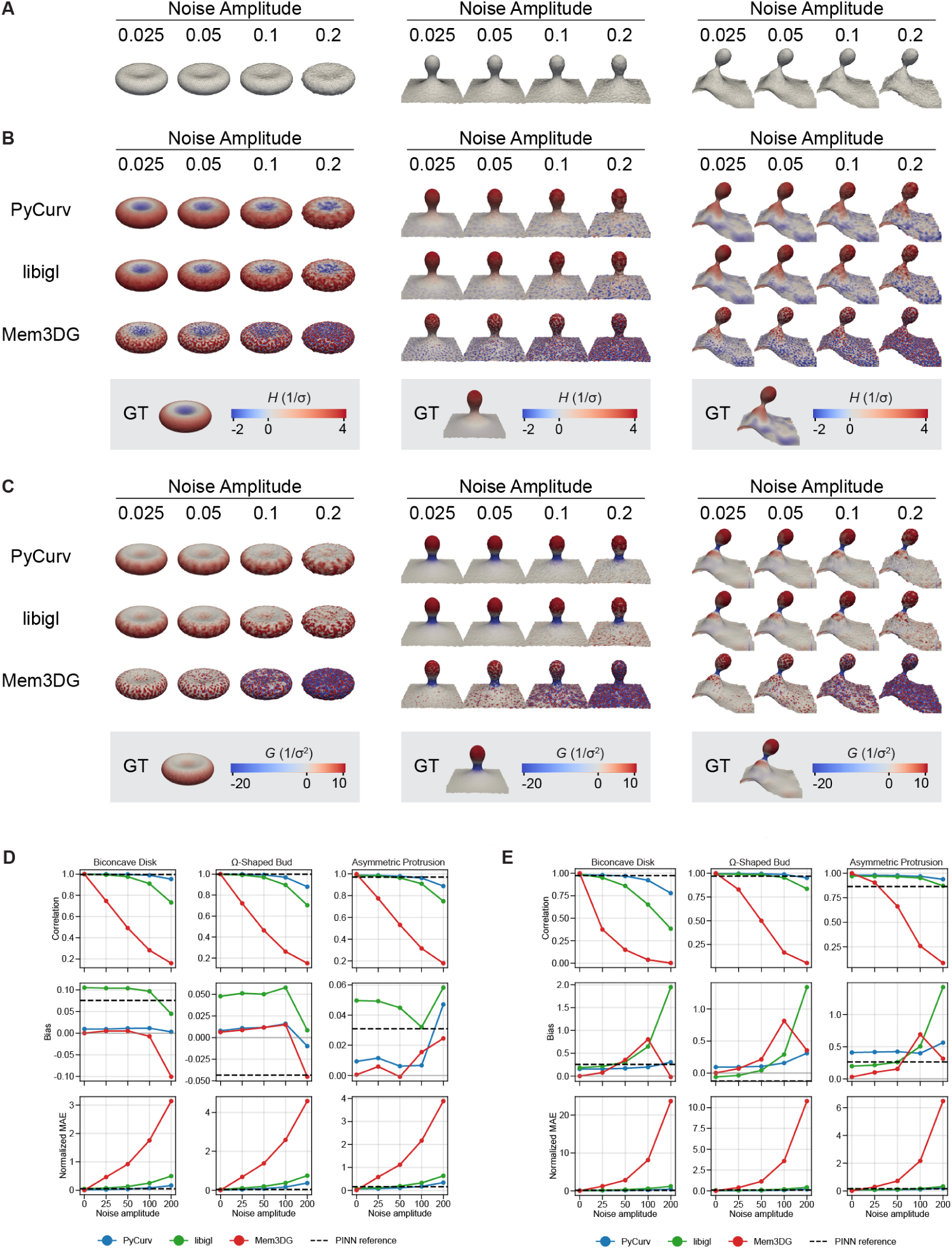
Curvature estimation using mesh-based membrane representations. (A) Synthetic membrane meshes with varying levels of noise applied to vertex positions. (B–C) Mean curvature (B) and Gaussian curvature (C) estimated using three mesh-based methods: PyCurv, libigl, and Mem3DG. (D–E) Accuracy of mean curvature (D) and Gaussian curvature (E) estimates evaluated against ground-truth values using Pearson correlation, mean signed error, and mean absolute error. Solid lines represent the results of mesh-based methods, while dashed lines indicate the corresponding PINN-based estimates for comparison.

**Figure S10:**
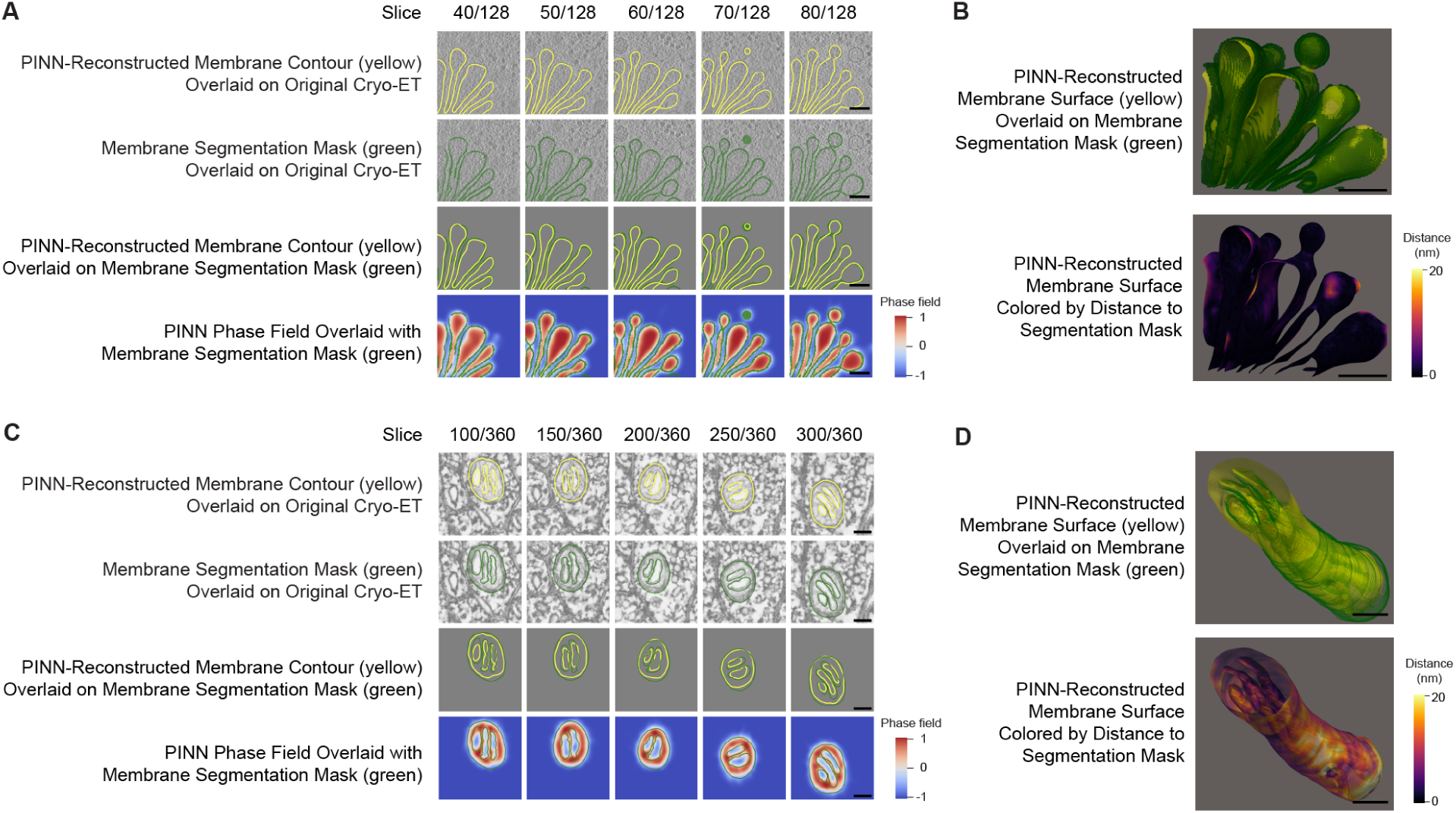
Comparison of PINN-reconstructed membrane geometries with reference segmentations. For the Golgi apparatus, we used a voxel-based segmentation generated using MemBrain-seg, whereas for the mitochondrion, we used manual segmentation defined on individual image slices. (A) and (C) Slice-by-slice comparison of PINN-reconstructed membrane contours (red), reference segmentations (green), original cryo-ET images, and reconstructed phase fields (color bar at the bottom right) for (A) the Golgi apparatus and (C) the mitochondrion. (B) and (D) Three-dimensional visualizations of (B) the Golgi apparatus and (D) the mitochondrion. Top: PINN-reconstructed membrane surfaces (yellow) overlaid with reference segmentations (green). Bottom: PINN-reconstructed surfaces colored by distance to the nearest reference segmentation point (color bar at the bottom right). Scale bars represent 100 nm in all panels.

**Table S1:** Quantitative evaluation of the geometric accuracy of PINN-reconstructed Golgi apparatus and mitochondrion membranes against the reference segmentation mask. Reconstruction accuracy was assessed using surface Dice at specified distance tolerances, average surface distance (ASD), and Hausdorff distance (HD).

| Metric | Golgi Apparatus | Mitochondrion | Unit |
|---|---|---|---|
| Surface Dice (tolerance = 5 nm) | 0.815 | 0.290 | - |
| Surface Dice (tolerance = 10 nm) | 0.966 | 0.612 | - |
| Surface Dice (tolerance = 15 nm) | 0.989 | 0.840 | - |
| Average Surface Distance | 3.02 | 9.10 | nm |
| 95th Percentile Hausdorff Distance | 10.61 | 20.4 | nm |
| Hausdorff Distance | 28.8 | 62.0 | nm |

**Figure S11:**
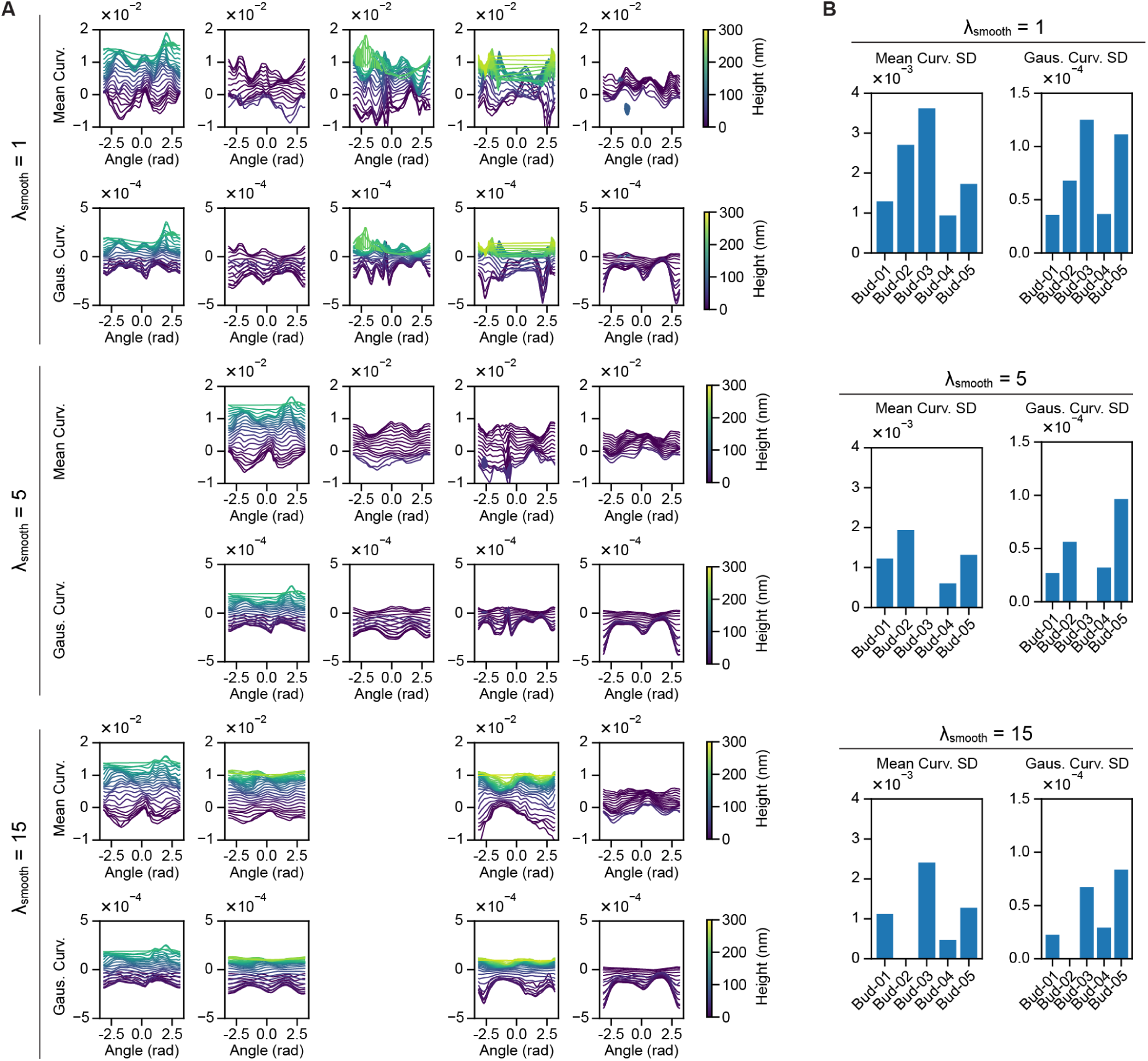
Curvature heterogeneity around the neck of endocytic pits is robust to variations in the smoothing loss weight. We performed PINN reconstructions with different smoothing loss weights, *λ*_smooth_ = 1, 5, 15, while keeping the other loss weights fixed, and evaluated the curvature distributions around the neck of endocytic pits. (A) Distributions of mean and Gaussian curvature along individual contour rings. (B) Standard deviations of mean and Gaussian curvature around the neck. Data are omitted for cases in which the PINN reconstruction failed.

